# BBB pathophysiology independent delivery of siRNA in traumatic brain injury

**DOI:** 10.1101/2020.06.26.173393

**Authors:** Wen Li, Jianhua Qiu, Xiang-Ling Li, Sezin Aday, Jingdong Zhang, Grace Conley, Jun Xu, Robert Langer, Rebekah Mannix, Jeffrey M. Karp, Nitin Joshi

## Abstract

Small interfering RNA (siRNA) represents a powerful strategy to mitigate the long-term sequelae of traumatic brain injury (TBI). However, poor permeability of siRNA across the blood brain barrier (BBB) poses a major hurdle. One approach to overcome this challenge involves treatment administration while the BBB is physically breached post-injury. However, this approach is only applicable to a subset of injuries with substantial BBB breach and can lead to variable therapeutic response due to the heterogeneity of physical breaching of BBB in TBI. Moreover, since physical breaching of BBB is transient, this approach offers a limited window for therapeutic intervention, which is not ideal as repeated dosing beyond the transient window of physically breached BBB might be required. Herein, we report a nanoparticle platform for BBB pathophysiology-independent delivery of siRNA in TBI. We achieved this by combined modulation of surface chemistry and coating density, which maximized the active transport of nanoparticles across BBB. Intravenous injection of engineered nanoparticles, within or outside the window of physically breached BBB in TBI mice resulted in 3-fold higher brain accumulation compared to conventional PEGylated nanoparticles and demonstrated up to 50% gene silencing. To our knowledge, this is the first reported example of BBB pathophysiology-independent drug delivery in TBI, and the first time combined modulation of surface chemistry and coating density has been shown to tune BBB penetration of nanoparticles. Taken together, our approach offers a clinically relevant approach to develop siRNA therapeutics for preventing long-term effects of TBI and deserves further exploration.

## Introduction

Referred to as a ‘silent epidemic’, traumatic brain injury (TBI) is a leading cause of death and disability among children and young adults, with millions of people sustaining TBI each year in accidents, sports, and military conflicts (*1–3*). TBI may lead to potentially long-lasting neurological dysfunctions, memory disturbances, behavioral changes, speech irregularities, and gait abnormalities. It has also been implicated in the development of neurodegenerative disease, particularly chronic traumatic encephalopathy, Alzheimer’s disease, and Parkinson’s disease (*4–7*).c (*8, 9*). Following primary injury, which is a result of mechanical impact to the brain., secondary injury gradually occurs as a consequence of destructive cellular and molecular events. This can ultimately lead to neuronal and glial cell death, tissue damage, and atrophy (8-10). While prevention strategies offer the best means to mitigate primary injury, strategies targeting secondary injury enable direct intervention by diminishing the cascade of destructive events. However, current treatments for TBI are limited to palliative care, and no effective strategies are available to interrupt the pathological cascade leading from TBI to neurodegeneration.

Nucleic acid-based therapeutics, especially small interfering RNA (siRNA) can efficiently and specifically silence the expression of target genes, including those that are traditionally considered to be “untreatable” by small molecule drugs (*11*). Thus, siRNA-based therapies can target specific pathological pathways to mitigate disease progression and offer a precision medicine approach for TBI treatment. Unfortunately, there are multiple biological barriers towards the delivery of siRNA into brain tissue; the blood brain barrier (BBB) being the most important one (*11–13*). One approach to overcome BBB in TBI involves systemic administration of therapeutics while the BBB is physically breached post-injury (*14–16*). However, physical breaching of BBB in TBI is highly heterogeneous and varies greatly among patients, depending on the extent of primary injury (*17–20*). Moreover, BBB can self-repair within a few hours to days post-injury to restore its integrity (*16–20*). Hence, physical breaching of BBB offers a limited window for therapeutic interventions, which is not ideal as the secondary injury can last months and may require repeated dosing beyond this limited time window. There is therefore an unmet need to develop approaches that allow BBB pathophysiology-independent delivery of siRNA in TBI.

Herein, we report a nanoparticle (NP) platform that enables brain delivery of siRNA, independent of BBB pathophysiology in TBI. We achieved this by combined modulation of surface coating chemistry and coating density on nanoparticles, which maximized their active transport through BBB. Incorporating certain surface coatings on NPs has been previously shown to augment their active penetration across BBB in multiple brain diseases (*21–24*). However, a surface coating that can facilitate active transport of siRNA NPs across BBB in TBI has not been reported previously. We first aimed to identify a surface coating and coating density that can maximize the active penetration of siRNA-loaded NPs across intact BBB. Specifically, we formulated NPs using poly(lactic-co-glycolic acid) (PLGA), a biodegradable and biocompatible polymer that exists in several FDA approved products (*25, 26*), and performed a systematic study to correlate the impact of different surface coatings and their coating densities on the penetration of PLGA NPs across intact BBB in healthy mice (Fig. 1A). Brain accumulation of NPs was found to be dependent on the type of surface coating and the coating density, and combined modulation of these two parameters maximized the penetration of NPs across BBB. Interestingly, surface coating and its coating density also impacted the uptake of siRNA-loaded NPs by neural cells and their gene silencing efficiency. Using an established mouse model of weight drop induced TBI (*27, 28*), we showed that the optimal NPs delivered siRNA to brain when administered during early or late injury periods, which correspond to physically breached BBB and intact, self-repaired BBB, respectively. Finally, intravenous administration of NPs loaded with siRNA against tau (a proof of concept target), during early or late injury period in TBI mice achieved up to 50% reduction in tau expression,. Overall, we demonstrate that BBB pathophysiology-independent delivery of siRNA in TBI is feasible and deserves further exploration.

**Fig. 1.**
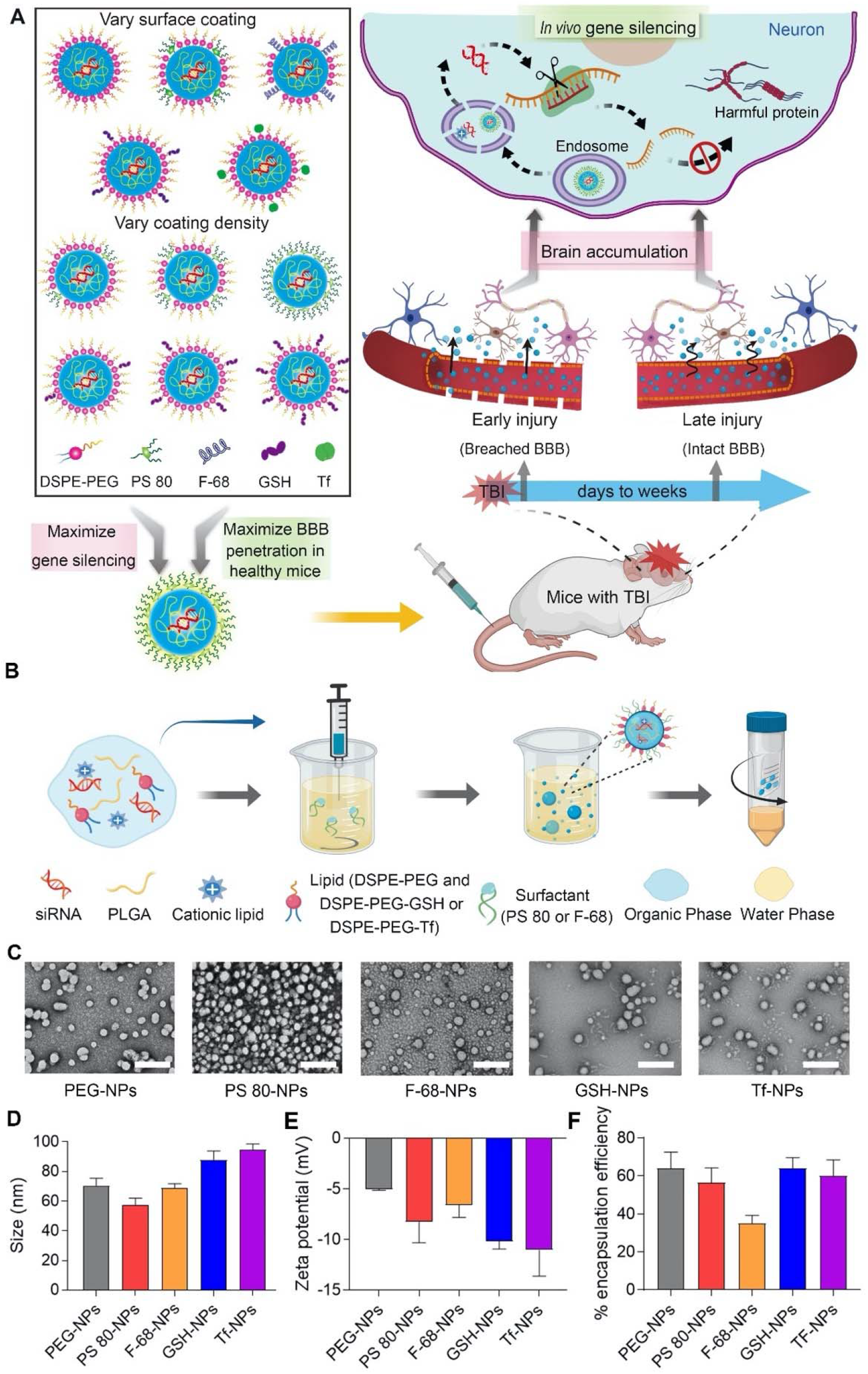
Nanoparticles (NPs) with different surface coatings were prepared to achieve BBB pathophysiology independent delivery of siRNA in TBI. (**A**) Schematic illustrating the overall study design. siRNA loaded NPs with different surface coating chemistries and coating densities were compared for their *in vitro* uptake and gene silencing efficiency in neural cells as well as their ability to cross intact BBB in healthy mice. NPs with maximum gene silencing efficiency and BBB permeability were then evaluated in TBI mice to determine brain accumulation and gene silencing efficiency when administered during early injury or late injury periods, corresponding to physically breached BBB and intact BBB, respectively. Upon neuronal uptake of NPs, siRNA is released and silences the harmful proteins involved in TBI pathophysiology. (**B**) Schematic for the preparation of siRNA-loaded poly(lactic-*co*-glycolic acid) (PLGA) NPs by a modified nanoprecipitation method. DSPE-PEG was used to impart stealth character. In addition, polysorbate 80 (PS 80), poloxamer 188 (F-68), DSPE-PEG-glutathione (GSH), or DSPE-PEG-transferrin (Tf) were used to augment BBB penetration. (**C**) Transmission electron microscopy (TEM) images of siRNA-loaded NPs having different surface coatings. Scale bar: 200 nm. (**D** and **E**) Size and zeta potential of siRNA-loaded NPs having different surface coatings, as analyzed by dynamic light scattering. (**F**) Encapsulation efficiency of siRNA in different NPs. Data in (D-F) are mean ± SD of technical repeats (*n* = 3, experiment performed at least twice).

## Results

### Sub 100 nm-sized siRNA NPs were prepared with different surface coatings

PLGA-based siRNA NPs with five different surface coatings were engineered to interact with BBB and neural cells by a variety of mechanisms. Polyethylene glycol (PEG) conjugated to 1,2-distearoyl-sn-glycero-3-phosphoethanolamine (DSPE) was incorporated in NPs for “stealth” effect to extend their systemic circulation time (*29*). In addition to DSPE-PEG, we also included one of the four different surface coatings that have previously shown to augment active penetration of NPs across BBB in other diseases. Specifically, we chose polysorbate 80 (PS 80), poloxamer 188 (Pluronic F-68), DSPE-PEG-glutathione (GSH), and DSPE-PEG-transferrin (Tf). PS 80 and F-68 have been previously shown to promote drug delivery across BBB by binding to endogenous apolipoproteins and interacting with lipoprotein receptors on BBB (*30–32*). GSH, a BBB shuttle peptide that has reached phase II clinical trials can transport NPs through BBB *via* GSH transporters (*33, 34*). Tf has been shown to facilitate brain delivery of NPs by binding to Tf receptors expressed on brain endothelium (*35, 36*).

siRNA-loaded NPs with different surface coatings were prepared using a modified nanoprecipitation method (Fig. 1B). Since NPs with small size (<100 nm) were previously demonstrated to exhibit more BBB penetration than larger ones (*37, 38*), we tuned the size of NPs in this study to <100 nm. Specifically, we tested different organic solvents for preparing NPs. The polarity of organic solvent influences the dispersion of polymer in aqueous solution, thus affecting the size of NPs. As shown by transmission electron microscopy (TEM) (Fig. 1C) and dynamic light scattering (DLS) analysis (Fig. 1D), spherical NPs with hydrodynamic diameters between 55-95 nm were formed with dimethylformamide (DMF), whereas larger NPs were obtained when using solvents with lower polarity, such as acetone and tetrahydrofuran (Fig. S1). Thus, DMF was chosen to formulate NPs for further studies. All NPs had a narrow size distribution, as suggested by the polydispersity index (Fig. S2), and had negative surface charge (Fig. 1E). The encapsulation efficiency of siRNA in different NPs was studied using Cy3 labeled scrambled siRNA. All NPs demonstrated high encapsulation efficiency (55-65%) except for NPs coated with F-68, which demonstrated a relatively lower encapsulation efficiency of 35% (Fig. 1F). Since F-68 impacted siRNA encapsulation, we did not use F-68 coated NPs for subsequent experiments. Finally, NPs showed slow and sustained release of siRNA in phosphate-buffered saline (PBS) (pH 7.4) at 37°C (Fig. S3).

### Surface coating impacts the uptake of NPs by neural cells and gene silencing efficiency

Efficient cellular uptake is the premise for achieving gene silencing. However, cell membrane is a barrier for siRNA delivery as negatively charged siRNA cannot cross the anionic cell membrane by itself. Therefore, we evaluated the uptake of siRNA-loaded NPs by Neuro-2a cells. NPs were loaded with scrambled siRNA labeled with a fluorescence probe (Dy677) to image cellular uptake by confocal laser scanning microscope (CLSM). Cells were incubated with free siRNA or siRNA-loaded NPs having different surface coatings at 37°C for 2 h, followed by imaging using CLSM. Cells incubated with free siRNA showed negligible fluorescence signal, which is expected. PEG-coated NPs (PEG-NPs) and Tf-coated NPs (Tf-NPs) also resulted in a weak fluorescence signal, suggesting that PEG and Tf do not facilitate the interaction between NPs and neural cells (Fig. 2A). Cells treated with GSH-coated NPs (GSH-NPs) or PS 80-coated NPs (PS 80-NPs) showed higher fluorescence signal compared to those treated with free siRNA, PEG-NPs or Tf-NPs, indicating that GSH and PS 80 facilitated the transport of siRNA-loaded NPs into neural cells. Particularly, PS 80-NPs exhibited the highest cellular uptake among all NPs, as suggested by an intense fluorescence signal in cells (Fig. 2A). We also performed flow cytometry to quantify cellular uptake of different NPs (Fig. S4). Consistent with CLSM analysis, the intensity of fluorescence signal in cells was found to be dependent on the surface coating chemistry of NPs. Cells treated with PS 80-NPs exhibited the strongest fluorescence signal, which was 20-fold higher than cells treated with PEG-NPs.

**Fig. 2.**
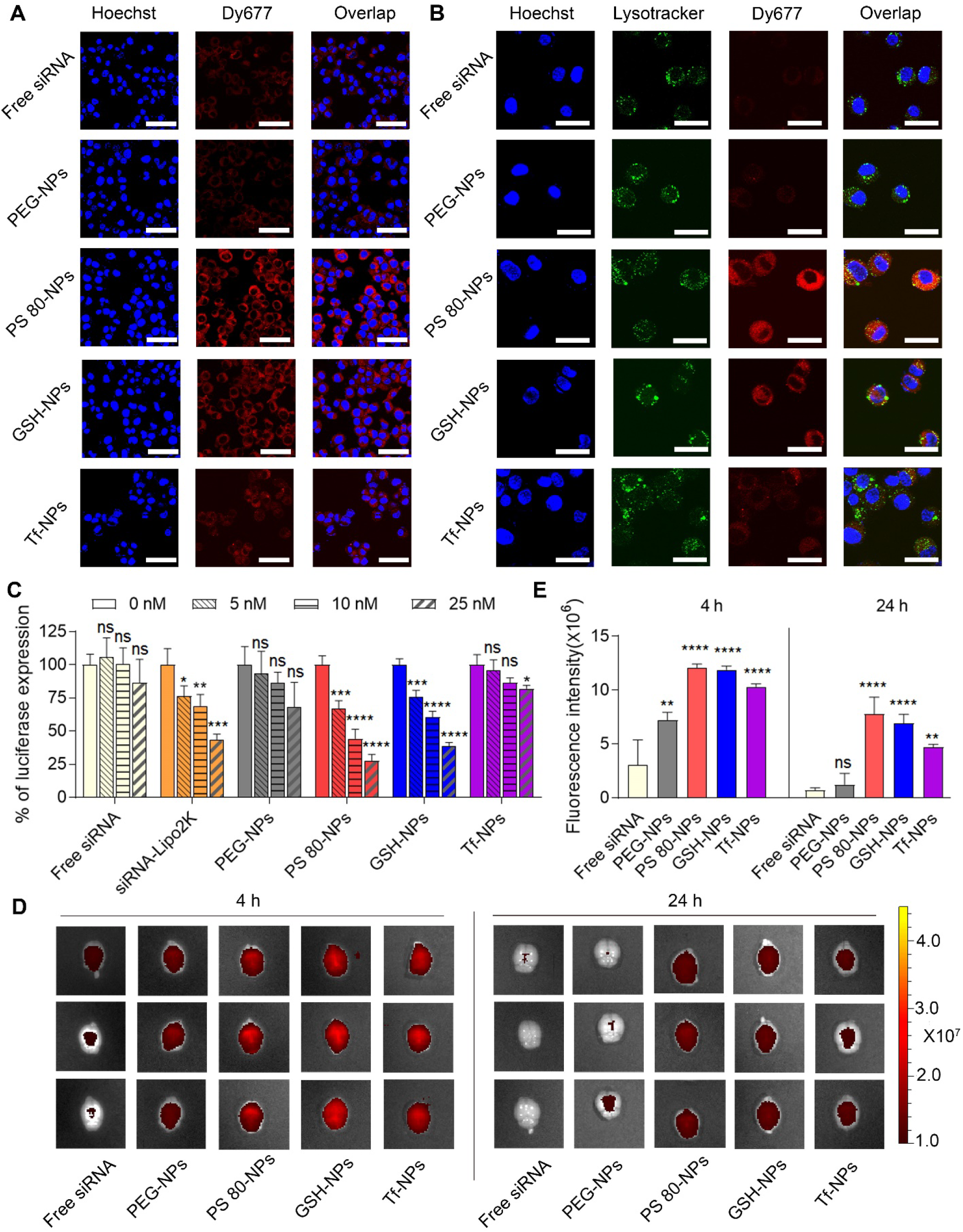
Surface coating impacts the uptake of siRNA-loaded NPs by neural cells, gene silencing efficiency *in vitro* as well as penetration of NPs across intact BBB. (**A**) Confocal laser scanning microscope (CLSM) images of Neuro-2a cells incubated with free siRNA or siRNA-loaded NPs having different surface coatings (PEG-NPs, PS 80-NPs, GSH-NPs or Tf-NPs) at 37 °C for 2 h. Dy677 labeled scrambled siRNA (red signal) was used. Nuclei were stained with Hoechst 33342 (blue signal). Scale bar: 50 μm. (**B**) CLSM images showing the endosomal escape of siRNA in Neuro-2a cells. Cells were incubated with free siRNA or siRNA-loaded NPs having different surface coatings at 37 °C for 2 h. Dy677 labeled siRNA (red signal) was used. Nuclei were stained by Hoechst 33342 (blue signal) and endosomes were stained with LysoTracker Green (green signal). Scale bar: 30 μm. (**C**) Luciferase expression in Neuro-2a cells. Luciferase-expressing Neuro-2a cells were incubated for 24 h with medium containing different concentrations of free luciferase siRNA (Free siRNA), luciferase siRNA-lipofectamine 2000 complex (siRNA-Lipo2K), or luciferase siRNA-loaded NPs having different surface coatings. Following an additional 48 h incubation with medium only, luciferase expression was quantified using a luminescence assay. \**P* < 0.05, \*\**P* < 0.01, \*\*\**P* < 0.001, \*\*\*\**P* < 0.0001 compared to 0 nM siRNA. (**D**) *In vivo* imaging system (IVIS) images of brains from three representative healthy mice, excised at 4 or 24 h after intravenous injection of free siRNA or siRNA-loaded NPs having different surface coatings (50 nmol siRNA/kg). Dy677 labeled scrambled siRNA was used. (**E**) Fluorescence intensity measured over excised mice brains using IVIS. \*\**P* < 0.01, \*\*\*\**P* < 0.0001 compared to free siRNA. Data in (C) are mean ± SD of technical repeats (*n* = 3, experiment performed at least twice). Data in (E) are mean ± SD (*n* = 3 mice/group, experiment performed twice). *P*-values were determined using one-way analysis of variance (ANOVA) with Tukey’s post-hoc analysis.

Following cellular uptake, siRNA must escape from endosomes to reach its target RNA in cytoplasm for gene silencing. Endosomal escape is one of the major obstacles for non-viral siRNA delivery, as siRNA trapped in endosomes will eventually undergo enzymatic digestion (*39*). To evaluate if siRNA could escape from endosomes, we used LysoTracker Green – a fluorescence probe to label endolysosomes and observed the co-localization of LysoTracker Green signal with signal from Dy677 labeled siRNA. After 2 h incubation of cells with NPs, siRNA was found to escape from endolysosomes into cytoplasm, as demonstrated by the separation of red (siRNA) and green (endolysosome) fluorescence signals in the cytoplasm (Fig. 2B).

We next sought to determine whether surface coating chemistry of NPs impact gene silencing efficiency. Luciferase-expressing Neuro-2a cells were incubated with varying concentrations of luciferase siRNA-loaded NPs having different surface coatings for 24 h, followed by incubation with medium only for an additional 48 h. Luciferase expression was quantified using a luminescence assay. Free siRNA and siRNA complexed with a commercially available transfection agent–lipofectamine 2000 (siRNA-Lipo2K) were used as controls. While cells treated with free siRNA did not result in any noticeable gene silencing, siRNA-Lipo2K silenced the luciferase gene in a dose-dependent manner (Fig. 2C). Gene silencing efficiency was found to be dependent on the surface coating chemistry of NPs. While PEG-NPs and Tf-NPs showed gene silencing only at the highest concentration of siRNA, PS 80-NPs and GSH-NPs resulted in significant gene silencing at all tested doses. In particular, PS 80-NPs exhibited the highest gene silencing, with 70% knockdown in luciferase expression at a concentration of 25 nM siRNA. Gene silencing efficiency of PS 80-NPs was also demonstrably greater than that mediated by Lipo2K. Notably, none of the NPs resulted in obvious decrease in the viability of cells compared to cells incubated with culture medium only (Fig. S5).

### Penetration of NPs across intact BBB depends upon surface coating

Next, we evaluated the brain accumulation of siRNA-loaded NPs having different surface coatings across intact BBB in healthy mice. Dy677 labeled scrambled siRNA was loaded into NPs. Free siRNA or siRNA loaded-NPs having different surface coatings were administered intravenously to healthy mice at siRNA dose of 50 nmol/kg. Mice were euthanized at 4 h and 24 h post-injection, and brains were harvested for imaging using *in vivo* imaging system (IVIS) (Fig. 2, D and E). Free siRNA and siRNA-loaded PEG-NPs showed weak fluorescence signals in the brain, suggesting poor penetration across BBB. PS 80-NPs, GSH-NPs, and Tf-NPs showed significantly higher fluorescence signals in the brain compared to PEG-NPs and free siRNA at both 4 h and 24 h time points. Impressively, brains from mice injected with PS 80-NPs exhibited the brightest fluorescence signal compared to mice injected with other formulations. Collectively, our data suggest that surface coating chemistry greatly impacts BBB penetration of NPs. For further studies, we chose to evaluate PS 80-NPs and GSH-NPs, due to their higher gene silencing efficiency and more effective penetration across intact BBB compared to PEG-NPs and Tf-NPs.

### Surface coating density impacts cellular uptake of NPs, gene silencing efficiency and penetration of NPs across intact BBB

We next modulated the surface coating density to understand its impact on the uptake of siRNA-loaded NPs by neural cells and the gene silencing efficiency. NPs coated with low (L), medium (M), or high (H) density of PS 80 or GSH were prepared. Quantitative values for surface coating densities are shown in Table S1. DLS analysis showed that the sizes of these NPs were all in the range of 50-90 nm (Fig. S6A). NPs became more negatively charged with an increase in the coating densities of PS 80 and GSH (Fig. S6B) with no impact on the encapsulation efficiency of siRNA (Fig. S6C). To study the impact of coating density on cellular uptake, Neuro-2a cells were incubated for 2 h with Dy677 labeled siRNA-loaded NPs having different coating densities of PS 80 or GSH, followed by imaging using CSLM. Cellular uptake of PS 80-NPs increased with an increase in the coating density of PS 80 (Fig. 3A). Flow cytometry showed 80-folds higher cellular uptake of PS 80 (H)-NPs compared to PEG-NPs (Fig. S7). Cellular uptake of GSH-NPs also increased with an increase in the coating density of GSH, but only modestly (Fig. 3A and Fig. S7). Coating density of PS 80 also impacted the gene silencing efficiency of NPs (Fig. 3B). Luciferase-expressing Neuro-2a cells were treated for 24 h with varying concentration of luciferase siRNA-loaded NPs having different coating densities of PS 80 or GSH. Following an additional 48 h incubation with medium only, luciferase expression was quantified. We observed dose-dependent decrease in the expression of luciferase for all the three PS 80-NPs having (L), (M), or (H) coating density. PS 80 (H)-NPs showed highest gene silencing, with 90% reduction of luciferase expression in Neuro-2a cells at a siRNA dose of 10 nM. Although coating density of GSH also impacted the gene silencing efficiency, the difference between different coating densities was not statistically significant. None of the NPs resulted in obvious decrease in the viability of cells compared to cells incubated with medium only (Fig. S8).

**Fig. 3.**
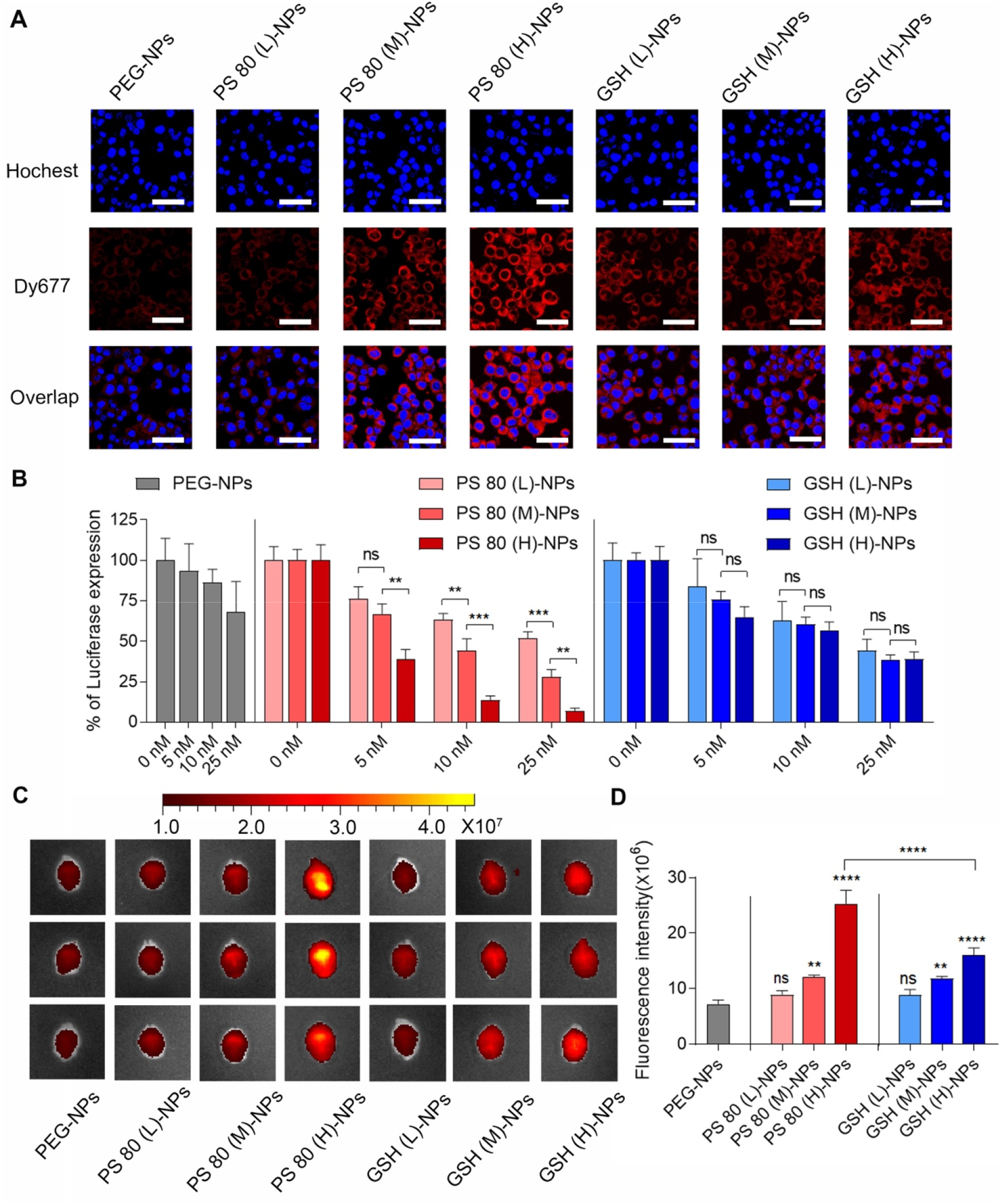
Surface coating density impacts the uptake of siRNA-loaded NPs by neural cells, gene silencing efficiency *in vitro* as well as penetration of NPs across intact BBB. (**A**) CLSM images of Neuro-2a cells incubated with siRNA-loaded NPs having different coating densities of PS 80 or GSH at 37 °C for 2 h. PEG-NPs were used as control. Dy677 labeled scrambled siRNA (red signal) was loaded into NPs. Nuclei were stained with Hoechst 33342 (blue signal). Scale bar: 50 μm. (**B**) Luciferase expression in Neuro-2a cells. Luciferase-expressing Neuro-2a cells were incubated for 24 h with medium containing luciferase siRNA-loaded NPs having different coating densities of PS 80 or GSH at varying concentrations of siRNA. Following an additional 48 h incubation with medium only, luciferase expression was quantified using a luminescence assay. PEG-NPs were used as control. \*\**P* < 0.01 and \*\*\**P* < 0.001. (**C**) IVIS images of brains from three representative healthy mice, excised at 4 h after intravenous injection of siRNA-loaded NPs (50 nmol siRNA/kg) having different coating densities of PS 80 or GSH. PEG-NPs were used as control. Dy677 labeled scrambled siRNA was loaded into NPs. (**D**) Fluorescence intensity measured over excised mice brains using IVIS. \*\**P* < 0.01 and \*\*\*\**P* < 0.0001 compared to PEG-NPs. \*\*\*\**P* < 0.0001 for PS 80 (H)-NPs *versus* GSH (H)-NPs. Data in (B) are mean ± SD of technical repeats (*n* = 3, experiment performed at least twice). Data in are mean ± SD (*n* = 3 mice/group, experiment performed twice). *P*-values were determined by one-way ANOVA with Tukey’s post-hoc analysis.

To evaluate the impact of coating density on the penetration of NPs across intact BBB, NPs having different coating densities of PS 80 or GSH were intravenously administrated to healthy mice at a siRNA dose of 50 nmol/kg. Dy677 labeled scrambled siRNA was loaded into NPs. Brains were collected for IVIS imaging at 4 h post-injection. Increase in the coating density of PS 80 or GSH increased the brain accumulation of NPs (Fig. 3, C and D). Mice injected with PS 80 (H)-NPs exhibited the strongest fluorescence signal, which was about 4 times higher than PEG-NPs. Increasing the coating density of GSH also enhanced the transport of NPs across BBB, but GSH (H)-NPs showed significantly less BBB penetration than the PS 80 (H)-NPs.

We also investigated the mechanism of BBB penetration by PS 80 NPs in an *in vitro* BBB model developed with mouse brain endothelial cell line, bEnd.3 (Fig. S9A). bEnd.3 cells were seeded on matrigel-coated filter inserts until tight junction formation, as confirmed by measuring the transendothelial electrical resistance (TEER). Dy677 labeled scrambled siRNA-loaded PEG-NPs or PS 80 (H)-NPs suspended in medium were added on the apical side. Following 4 h incubation, aliquots from the basolateral side of the inserts were collected, and the amount of NPs that crossed the cell layer into the basolateral compartment was quantified by fluorescence. Compared to PEG-NPs, PS 80 (H)-NPs showed significantly higher penetration across the cell layer (Fig. S9B), which is consistent with *in vivo* brain accumulation results. However, when the same experiment was performed in the absence of serum in the culture medium, penetration of PS 80 (H)-NPs reduced significantly (Fig. S9C), which suggests that certain serum proteins indeed play a crucial role in the transport of PS 80 (H)-NPs across BBB. It has been reported that PS 80 can adsorb endogenous lipoprotein, such as ApoE or ApoA-I to facilitate the transport of NPs across BBB *via* lipoprotein receptor-mediated transcytosis (*30–32*). To confirm if lipoprotein plays a role in mediating the penetration of PS 80 (H)-NPs across BBB, we blocked the lipoprotein receptor (LRP1) by anti-LRP1 antibody, which decreased the permeability of PS 80 (H)-NPs (Fig. S9C), confirming the role of lipoprotein in PS 80-induced penetration of NPs across BBB.

*In vivo* circulation profiles of different siRNA formulations were also compared (Fig. S10). A longer circulation time allows NPs to have a greater tendency to accumulate in the target organ. Dy677 labeled free siRNA or Dy677 labeled siRNA-loaded PEG-NPs or PS 80 (H)-NPs were injected into mice *via* tail vein at siRNA dose of 50 nmol/kg. Blood was withdrawn at the indicated time points for the measurement of fluorescence signal. Free siRNA was eliminated rapidly from the blood, and was hardly detected 30 min post-injection. PEG-NPs significantly prolonged the circulation time of siRNA in blood, which could be explained by the protection and stealth effect of the PEG layer (*40, 41*). Interestingly, PS 80 (H)-NPs, which did not have the protective PEG layer on the surface, demonstrated similar circulation profile as PEG-NPs. This long circulation time is presumably attributable to the hydrophilicity and non-ionic structure of PS 80 that reduces the uptake of NPs by the reticuloendothelial system (*42*). Since PS 80 (H)-NPs showed maximum *in vitro* gene silencing efficiency, maximum penetration across intact BBB, and long circulation time in blood, we decided to evaluate them further in a mouse model of TBI.

### PS 80 (H)-NPs exhibit BBB pathophysiology-independent delivery of siRNA in a mouse model of TBI

Based on the combined modulation of surface coating chemistry and coating density, PS 80 (H)-NPs were selected as optimal NPs for further evaluation in TBI mice. We used the weight drop induced TBI model (Fig. 4A), which is a clinically relevant murine model of TBI (*27, 28*). Briefly, mice were anesthetized and subjected to head impact by dropping a 54 g weight from 60-inch height. We first characterized the time window of physically breached BBB following TBI using Evans blue (EB) penetration assay. EB dye binds to serum albumin in the bloodstream and does not cross the intact BBB under normal physiological conditions, but permeates physically breached BBB (*43*). Healthy mice or mice with TBI were injected with EB *via* tail vein. In the case of TBI mice, EB was injected at different time points post-injury. Two hours after EB administration, animals were perfused extensively with saline, and EB content in brain tissue was quantified. EB dye injected to TBI mice at 6 h and 24 h post-injury resulted in significantly higher EB content compared to healthy mice (Fig. 4B). EB dye injected 1 week post-injury and later resulted in substantially less brain accumulation compared to 24 h, indicating a decrease in EB permeability. Collectively, our data suggest that weight drop induced TBI resulted in physical breaching of BBB which increased over time, until at least 24 h post-injury. BBB was significantly repaired by 1 week, with complete recovery at 2 weeks post-injury (Fig. 4B).

**Fig. 4.**
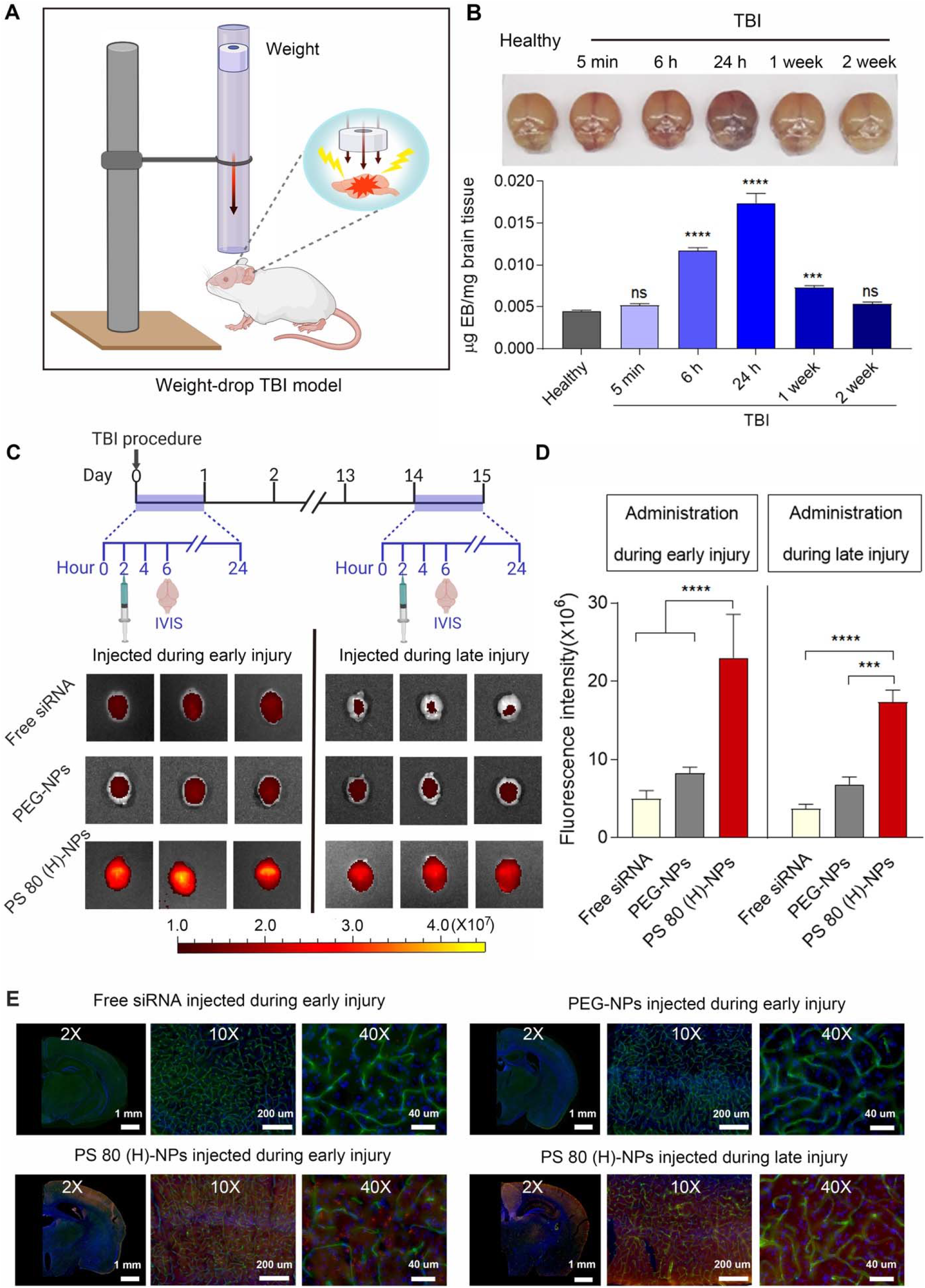
PS 80 (H)-NPs exhibit BBB pathophysiology-independent delivery of siRNA in a mouse model of TBI. (**A**) Schematic showing the experimental procedure for weight drop-induced TBI model. (**B**) Time window of physically breached BBB following TBI was characterized by Evans blue (EB) penetration assay. EB was intravenously injected into healthy mice or mice with TBI at different time points post-injury. Two hours after EB administration, brains were excised and photographed. EB content in brain tissue was also quantified and expressed as μg of EB per mg of brain tissue. \*\*\**P* < 0.001, \*\*\*\**P* < 0.0001 compared to healthy mice. (**C**) Experimental outline: Free siRNA, siRNA-loaded PEG-NPs, or PS 80 (H)-NPs were intravenously injected into mice after 2 h or 2 weeks of TBI procedure, corresponding to early and late injury, respectively. Four hours after the administration of siRNA formulations, brains were excised and imaged using IVIS. Dy677 labeled scrambled siRNA was used at 50 nmol/kg dose. IVIS images of brains from three representative healthy mice injected during early or late injury are shown. (**D**) Fluorescence intensity measured over excised mice brains using IVIS. \*\*\**P* < 0.001, \*\*\*\**P* < 0.0001. (**E**) Representative fluorescence microscopy images of brain sections from mice injected intravenously with free siRNA, siRNA-loaded PEG-NPs or PS 80 (H)-NPs during early or late injury. Dy677 labeled scrambled siRNA (red signal) was used. Nuclei were stained with Hoechst 33342 (blue signal), and blood vessels were labeled with FITC-lectin (green signal) Data in (B and D) are mean ± SD (*n* = 3 mice/group, experiment performed twice). *P*-values were determined by one-way ANOVA with Tukey’s post-hoc analysis.

Next, we investigated the penetration of Dy677 labeled siRNA-loaded PS 80 (H)-NPs in the weight drop induced TBI model (Fig. 4C). Free siRNA and siRNA-loaded PEG-NPs were used as control. To characterize penetration across physically breached BBB during early injury, siRNA-loaded NPs or free siRNA at 50 nmol/kg dose of scrambled siRNA were injected at 2 h post-injury. Brains were collected for IVIS imaging at 4 h post-injection (Fig. 4C). Weak fluorescence signal was detected in the brains of mice injected with free siRNA or siRNA-loaded PEG-NPs, indicating low brain accumulation (Fig. 4, C and D). By contrast, we observed a strong fluorescence signal in the brains of mice injected with siRNA-loaded PS 80 (H)-NPs, with a 5-fold and 3-fold higher fluorescence intensity when compared with free siRNA and siRNA-loaded PEG-NP groups, respectively (Fig. 4, C and D). To evaluate siRNA delivery across intact BBB during late injury, free siRNA or siRNA-loaded NPs were administrated intravenously 2 weeks post-injury, when BBB was self-repaired (Fig. 4C). Again, mice administered with siRNA-loaded PS 80 (H) NPs displayed the highest fluorescence signal in the brain (Fig. 4, C and D). These results indicate that PS 80 (H)-NPs can efficiently deliver siRNA into the brain, when administered during early or late injury period after TBI. This holds great potential for downregulating pathological targets involved in the secondary brain injury, a process that usually lasts weeks to months. To specifically localize the delivered siRNA within brain tissue, blood vessels were labeled with FITC-lectin by injecting it into mice *via* tail vein 10 min before euthanasia. Brains from mice were sectioned and observed with fluorescence microscope. There was nearly no detectable siRNA fluorescence signal in the brain of mice injected with free siRNA or siRNA-loaded PEG-NPs (Fig. 4E). However, in the brain sections from mice treated with siRNA-loaded PS 80 (H)-NPs, we observed substantial red fluorescence signal from Dy677 labeled siRNA in the cortex (Fig. 4E). Notably, we observed minimum co-localization of red fluorescence signal from Dy677 labeled siRNA with green fluorescence signal from FITC lectin-stained brain vessels. This indicates that most NPs accumulated in the extravascular brain tissue, which confirms the ability of PS 80 (H)-NPs to cross brain vasculature. Similarly, NPs administered at 2 weeks post-injury also showed penetration into the extravascular brain tissue.

### Tau siRNA loaded PS 80 (H)-NPs suppress tau expression in murine primary neuronal cells

Among various harmful pathways, tau pathology has been found to be highly associated with chronic neuroinflammation, neurodegeneration, and cognitive impairment caused by TBI (*44–46*). Tau is a microtubule-associated protein found mostly in neurons. Following TBI, tau becomes hyperphosphorylated and dissociates from microtubules, forming abnormal aggregates that are strongly linked to TBI-associated chronic traumatic encephalopathy and Alzheimer’s disease (*27*, *44–46*). Prior studies showed that the reduction of tau protein attenuated its aggregation by decreasing the amount of tau available for phosphorylation (*46–48*), representing potential interventions for TBI treatment. We therefore sought to investigate whether PS 80 (H)-NPs could efficiently deliver tau siRNA into the brains of TBI mice to suppress tau expression.

Tau silencing ability of tau siRNA-loaded PS 80 (H)-NPs was first evaluated *in vitro* in primary neurons of mice. Unlike cell lines, primary neurons maintain certain phenotypic features and functions of nervous systems, thereby representing a more relevant *in vitro* model for neuroscience research. The primary cortical neurons were prepared from freshly isolated mouse embryonic cortical tissues. After 10 days of culture, primary neurons displayed a high degree of neurite outgrowth and branching (Fig. 5A). At that point, neurons were treated for 24 h with medium only or medium containing free tau siRNA, tau siRNA-loaded PEG-NPs, tau siRNA-loaded PS 80 (H) NPs, or scrambled (control) siRNA-loaded PS 80 (H)-NPs at 15 nM concentration of siRNA. Following an additional 48 h incubation with medium only, western blot analysis was performed to measure the expression of tau in primary neurons. Tau siRNA-loaded PS 80 (H)-NPs dramatically downregulated tau expression in neurons with approximately 70% decrease of tau expression (Fig. 5B). In contrast, treatment with free tau siRNA, tau siRNA-loaded PEG-NPs, or control siRNA-loaded PS 80 (H)-NPs resulted in negligible reduction of tau expression (Fig. 5B). Tau knockdown by tau siRNA-loaded PS 80 (H) NPs was also shown to be dose-dependent (Fig. 5C). Inhibition of tau was further confirmed by immunofluorescence imaging (Fig. 5D). Compared with all the other groups, cells treated with tau siRNA-loaded PS 80 (H)-NPs showed much weaker green fluorescence, suggesting lower expression levels of tau protein.

**Fig. 5.**
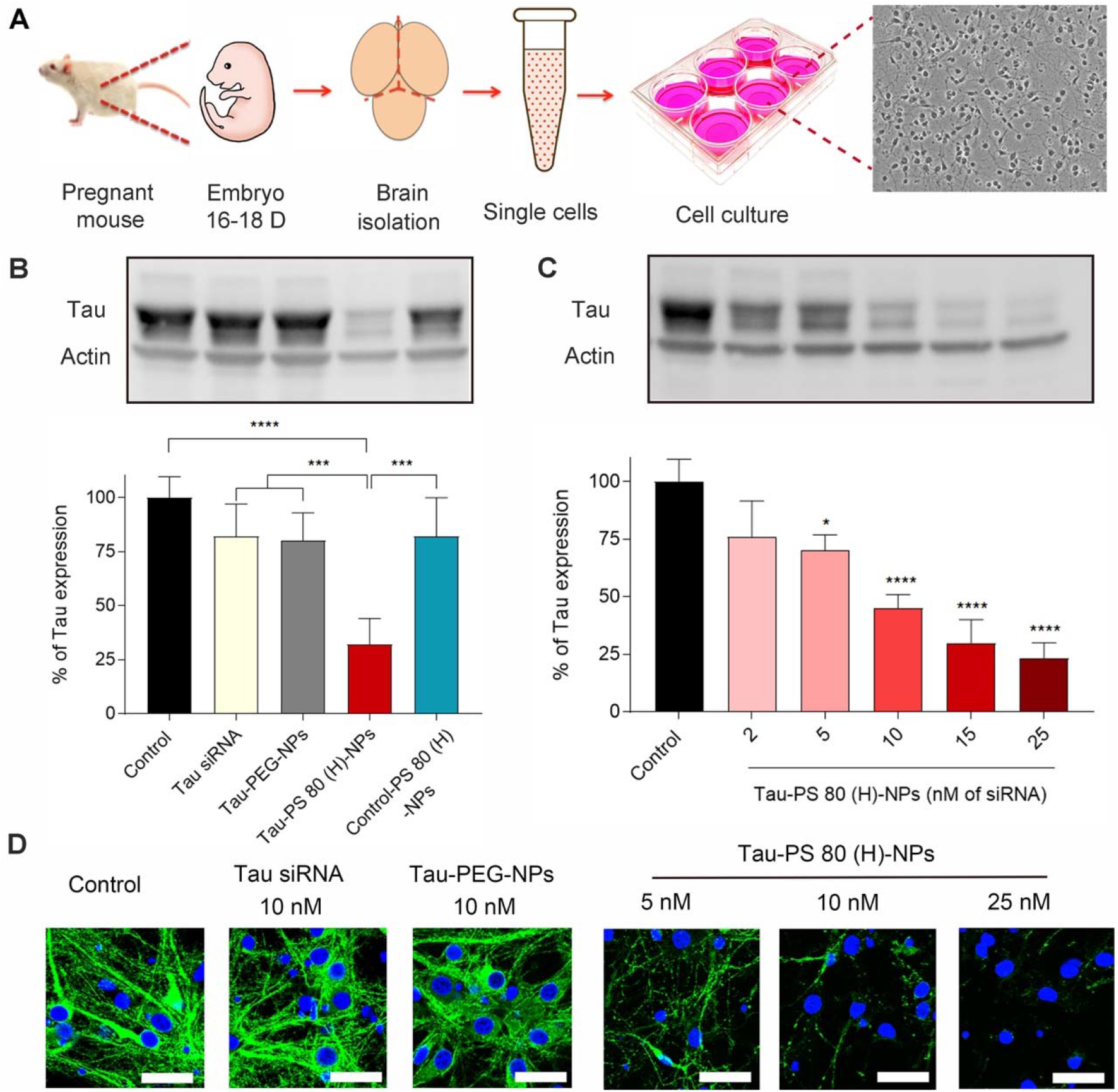
Tau siRNA-loaded PS 80 (H)-NPs suppress tau expression in murine primary neuronal cells: (**A**) Schematic illustration depicting the isolation of primary neuronal cells from mouse embryos, and bright-field image of primary cells after 10 days in culture. Pregnant female C57BL/6 mice at E18 were euthanized, the cortex was dissected from embryo brains and enzymatically digested to obtain single cell suspension. Finally, cells were cultured on plates pre-coated with poly-L-lysine. (**B**) Western blots and quantification of tau expression in primary neuronal cells. Cells were incubated for 24 h with only medium (control) or medium containing free tau siRNA (Tau siRNA), tau siRNA-loaded PEG NPs (Tau-PEG-NPs), tau siRNA-loaded PS 80 (H) NPs (Tau-PS 80 (H)-NPs), or scrambled siRNA-loaded PS 80 (H) NPs (Control-PS 80 (H)-NPs). The dose of siRNA was 15 nM. Following an additional 48 h incubation with medium only, western blot analysis was performed. \*\*\**P* < 0.001, \*\*\*\**P* < 0.0001. (**C**) Western blot and quantification of tau expression in primary neuronal cells treated with tau siRNA-loaded PS 80 (H)-NPs at different concentrations of siRNA. \**P* < 0.05, \*\*\*\**P* < 0.0001 compared to control. (**D**) Immunofluorescence images of tau expression in primary neuronal cells. Cell were treated for 24 h with medium only (control) or medium containing free tau siRNA (10 nM siRNA), tau-PEG-NPs (10 nM siRNA), or tau-PS 80 (H)-NPs (5, 10 or 25 nM siRNA). Following an additional 48 h incubation with medium only, immunofluorescence staining was performed. Nuclei were stained with Hoechst 33342 (blue signal) and tau was stained with the anti-tau primary antibody followed by Alexa Fluor® 488-labeled secondary antibody (green signal). Scale bar: 30 μm. Data in (B and C) are mean ± SD of technical repeats (*n* = 3, experiment performed twice). *P*-values were determined by one-way ANOVA with Tukey’s post-hoc analysis.

### Tau siRNA loaded PS 80 (H)-NPs silence tau expression during early and late injury phases with no systemic toxicity

We next investigated the tau silencing ability of tau siRNA-loaded PS 80 (H)-NPs in the weight drop induced model of TBI (Fig. 6, A and B). To investigate tau silencing during early injury phase, mice were intravenously administrated with PBS, free tau siRNA, or PS 80 (H)-NPs loaded with either scrambled (control) siRNA or tau siRNA at siRNA dose of 75 nmol/kg/day. Treatments were administered at 2 h and 1 day post-injury. On day 4, brains were harvested, and the cortex was isolated and processed for analysis of tau expression using western blotting. Free siRNA treatment resulted in negligible tau knockdown, whereas tau siRNA-loaded PS 80 (H)-NPs dramatically reduced the expression of tau by about 50% (Fig. 6A). Scrambled siRNA-loaded PS 80 (H)-NPs did not reduce tau expression. Interestingly, even when administered during the late injury period, i.e. 2 weeks after the injury, tau siRNA-loaded PS 80 (H)-NPs could block around 40% tau expression in the cortex (Fig. 6B). Immunohistochemical (IHC) staining analysis further confirmed the robust tau silencing ability of tau siRNA-loaded PS 80 (H)-NPs, when administered during early or late injury period (Fig. 6C).

**Fig. 6.**
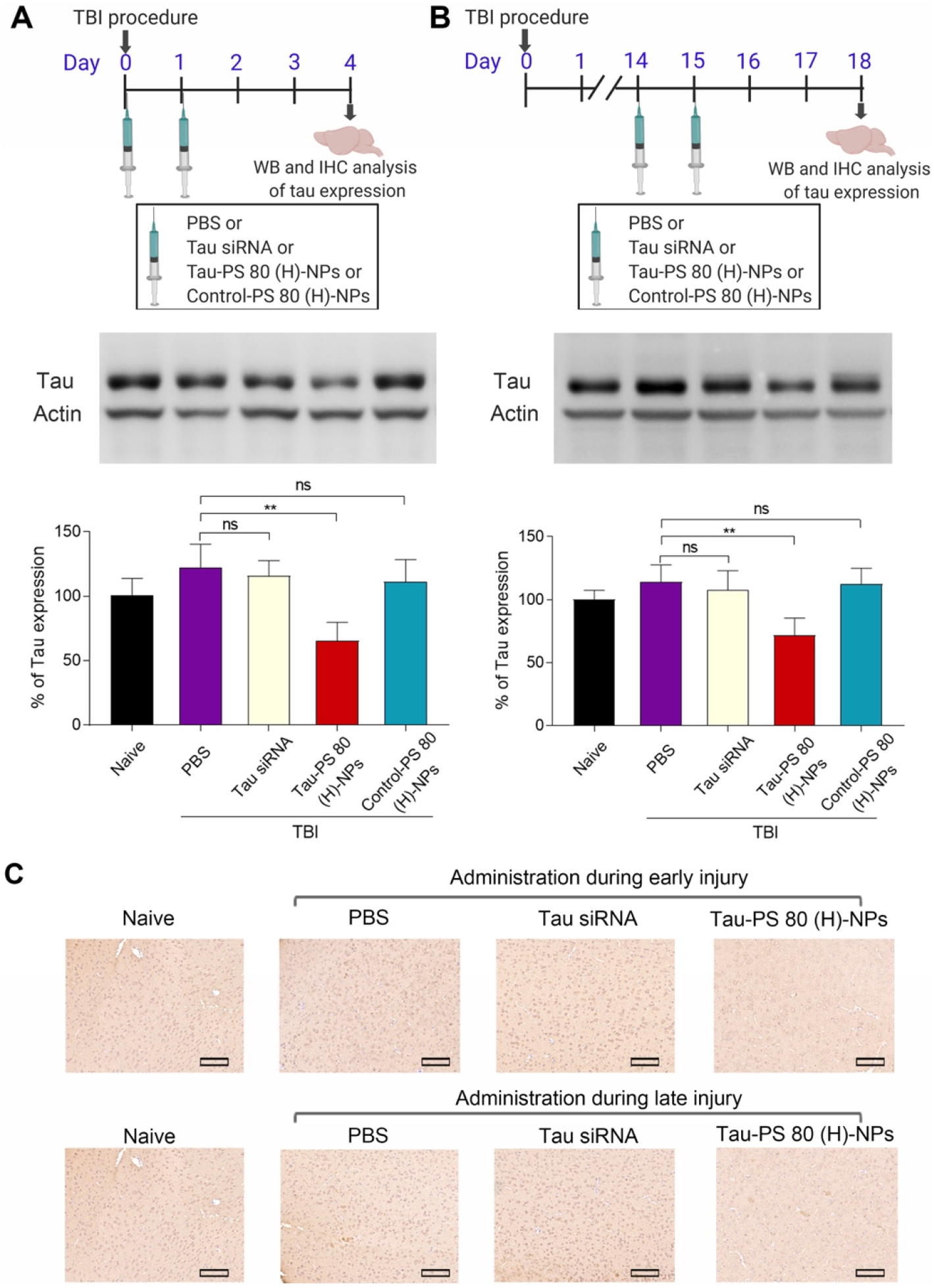
Tau siRNA loaded PS 80 (H)-NPs silence tau expression *in vivo* during early and late injury. (**A**) Experimental outline: To evaluate tau silencing efficiency during early injury, mice received tail vein injection with PBS, free tau siRNA (tau siRNA), tau siRNA-loaded PS 80 (H) NPs (tau-PS 80 (H)-NPs), or scrambled siRNA-loaded PS 80 (H)-NPs (control-PS 80 (H)-NPs) at 2 h and 1 day post-injury. siRNA dose was 75 nmol/kg/day. Brains were dissected to isolate cortex on day 4 for quantification of tau expression using western blot. Representative western blots and quantification of tau expression are shown. Naive animals were healthy mice with no treatment. \*\**P* < 0.01. (**B**) Experimental outline: To evaluate tau silencing efficiency during late injury, mice received tail vein injection of PBS, tau siRNA, tau-PS 80 (H)-NPs, or control-PS 80 (H)-NPs at day 14 and 15 post-injury. siRNA dose was 75 nmol/kg/day. Brains were harvested on day 18 for quantification of tau expression using western blot. Representative western blot analysis and quantification of tau expression were shown. \*\**P* < 0.01. (**C**) Immunohistochemical staining of tau expression in brain tissue sections of naive mice (healthy mice with no treatment) or TBI mice treated with PBS, tau siRNA, or tau-PS 80 (H)-NPs. Treatments were performed during the early or late injury phase at 75 nmol/kg/day dose of siRNA. Scale bar = 150 μm. Data in (A and B) are mean ± SD (*n* = 3 mice/group, experiment performed twice). *P*-values were determined by one-way ANOVA with Tukey’s post-hoc analysis.

We also evaluated the *in vivo* safety profile and systemic toxicity of PS 80 (H)-NPs. Healthy mice were injected with PBS or PS 80 (H)-NPs loaded with scrambled (control) siRNA or tau siRNA at a siRNA dose of 75 nmol/kg/day for 2 consecutive days. None of the treatment groups showed significant bodyweight reduction over 2 weeks after injection (Fig. S11). Additionally, hematoxylin and eosin (H&E) staining did not show any notable pathological changes in major organs, including lung, heart, liver, spleen, and kidney after 3 days of injection (Fig. S12). Hematology markers for mice that received PS 80 (H)-NPs loaded with tau siRNA or scrambled siRNA were within normal range and not significantly different from the PBS injected group at 3 days and 2 weeks after the last injection (Fig. S13). Finally, multiple blood biochemical parameters, including alkaline phosphatase (ALP), alanine aminotransferase (ALT), aspartate aminotransferase (AST), creatinine, and blood urea nitrogen (BUN) were evaluated for assessing effects on liver and kidney (Fig. S13). All biochemical parameters for mice injected with siRNA-loaded NPs were not significantly different from PBS injected mice. These results collectively indicate the safety of PS 80 (H)-NPs as siRNA nanocarriers.

## Discussion

To date, more than 30 clinical trials have investigated treatment options for TBI; however, none have successfully proceeded to phase III trials (*49*). siRNA represents a powerful therapeutic tool against TBI, yet the utility of siRNA for TBI treatment is hindered by multiple barriers, especially due to its inability to cross BBB. Direct injection of siRNA into brain tissue has been explored to bypass BBB (*50*), and few other efforts focused on the use of chemical/biological agents or physical stimuli to artificially induce the opening of BBB for siRNA transport. Although effective, these methods are either invasive or non-selective, which may add further complications to the injury. Recently, several NP platforms have been explored to deliver therapeutics across BBB for TBI treatment (*14–16*). However, these NPs usually rely on injury-induced physical breaching of BBB for brain accumulation, which may lead to highly variable therapeutic response due to the heterogeneous nature of physical breaching of BBB in TBI. Moreover, since physical breaching of BBB is transient, these approaches necessitate administration of the treatment within an extremely narrow time window after TBI, which is difficult to achieve in practice and has been an important contributing factor in the failure of several previous clinical trials (*50, 51*). Also, since secondary injury last months, repeated treatments beyond the transient window of physically breached BBB might be required. Hence, NPs that do not rely on BBB pathophysiology for brain accumulation are highly desired for reliable and long-term treatment of TBI.

This is the first reported example of BBB pathophysiology-independent delivery of siRNA loaded NPs in TBI. We achieved this by combined modulation of surface coating chemistry and coating density, which maximized the active transport of NPs across BBB. While modulation of surface coating chemistry alone has been previously shown to impact BBB penetration of nanoparticles, this is the first report demonstrating that combined modulation of surface coating chemistry along with coating density can further augment the active penetration of NPs through BBB. PS 80 (H)-NPs by far showed the best neural cell uptake, gene silencing efficiency, and brain accumulation in TBI mice when administered during early or late injury periods. Intravenous injection of PS 80 (H)-NPs, within or outside the window of physically breached BBB in TBI mice resulted in 3-fold higher brain accumulation compared to conventional PEGylated NPs. Interestingly, fluorescent microscopy images of brain sections in our study revealed that PS 80 (H)-NPs accumulate primarily in the cortex, which has been previously shown to play a crucial role in TBI pathogenesis (*49*). Although the mechanism of this preferential distribution is unknown at this time, the topic could potentially shed light on improving therapeutic targeting for TBI and merits more exploration in future studies.

PS 80 is a non-ionic surfactant that has been widely used in pharmaceutical and food industries. PS 80-coated NPs have been previously employed to promote drug delivery into brain for multiple applications, including glioblastoma (*32, 42*). However, it is difficult to directly compare brain accumulation observed in our study with literature due to the differences in particle type, drug type, and disease/animal model (*21*). Nevertheless, to our knowledge, the use of PS 80-coated NPs for brain delivery of siRNA in TBI treatment has never been explored previously. Additionally, our study is the first to demonstrate that PS 80 coating density is a critical parameter to maximize penetration of PS 80-coated NPs across BBB.

Compared to protein or peptide-based approaches to augment BBB penetration, PS 80 possesses the features of low cost, high stability, and low immunogenicity. PS 80-NPs developed here also demonstrated small size (about 60 nm), high siRNA encapsulation efficiency, and long circulation profile, comparable to PEG-NPs. Moreover, they were prepared by a robust one-step method, which greatly enhances the platform’s potential for large scale manufacturing and clinical translation. These features collectively suggest that PS 80 could serve as a promising tool for brain delivery of siRNA for TBI treatment.

As a proof-of-concept, we investigated the efficacy of PS 80 (H)-NPs to silence tau expression after TBI. Tau pathology is strongly linked to the TBI-caused neurodegeneration and brain dysfunction (*27, 44–46*). Therefore, therapies that reduce tau level or inhibit the phosphorylation or aggregation of tau represent potential interventions for TBI treatment. Although a few small molecule drugs have been reported to reduce tau level (*52,53*), they can potentially interact with other unintended and unknown targets. siRNA is a promising strategy to modulate tau levels due to its high efficacy and specificity. siRNA-mediated reduction of tau protein has been shown to attenuate tau aggregation and neuronal death by decreasing the amount of tau available for phosphorylation (*47, 48*). However, to our knowledge, systemic delivery of tau siRNA has not been investigated for the treatment of TBI. We showed that tau siRNA-loaded PS 80 (H)-NPs could not only dramatically knockdown tau expression in cultured primary neural cells, but also achieved 40-50% tau silencing in TBI mice, when administered within or outside the window of physically breached BBB, which correspond to early and late injury, respectively. This opens up exciting possibilities in TBI research by enabling long-term, non-invasive mitigation of harmful pathways.

There are some limitations in our study. Firstly, although we clearly demonstrated the significance of surface coating density to improve BBB penetration of NPs, we have limited our experiments to three different coating densities of PS80 and GSH. Future studies could explore a wider range of coating densities to further establish the extent to which BBB penetration can be enhanced. Secondly, we only proceeded with PS 80 (H)-NP formulation for evaluation in TBI mice. We acknowledge that GSH-NPs also showed substantial accumulation across intact BBB and merits more exploration. Finally, due to the complexity of TBI pathobiology, there is as of yet a limited understanding of the pathways that lead to neurobehavioral sequelae of TBI. One of these pathways almost certainly involves tau protein, which we have chosen as the target for our proof-of-concept study. However, suppressing tau pathology alone may not be sufficient to result in a significant change in functional outcomes. For this reason, we have not included functional or behavior evaluations in this study. Future studies could explore siRNA against additional pathobiological pathways, such as cell apoptosis, neuroinflammation, edema, calcium overload, or coagulation, as well as larger animal models of TBI (such as rabbit or pig), and evaluate more clinically relevant therapeutic benefits, including functional outcomes and performance in behavioral tasks. Moreover, since siRNAs against other targets could be readily incorporated into this NP platform, it also presents a potential research tool for the validation and modulation of novel therapeutic targets for neurological diseases, particularly those which are considered as “undruggable” targets.

In conclusion, our work elucidates a clinically relevant approach for developing siRNA therapeutics to prevent the long-term effects of TBI. To our knowledge, this is the first reported example of BBB pathophysiology-independent siRNA delivery in TBI, and for the first time we demonstrate that combined modulation of surface chemistry and coating density can be an efficient tool to tune BBB penetration of nanoparticles. Our novel NP approach also increases the potential of siRNA as a therapeutic tool against other neurological diseases, such as Alzheimer’s and Parkinson’s diseases, and therefore holds great promise for the design and translation of precision therapeutics for brain disorders.

## Materials and Methods

### Synthesis of DSPE-PEG-GSH and DSPE-PEG-Tf

Transferrin (Sigma Aldrich) was first thiolated using 2-iminothiolane (Traut’s reagent) as previously described (*54*). Thiolated Tf was then immediately used for conjugation with DSPE-PEG-maleimide with PEG molecular weight 3400 (DSPE-PEG-MAL) (NANOCS) *via* thiol-Michael addition reaction (*54, 55*). DSPE-PEG-Mal was first dissolved in dry DMF, and then 10-fold molar excess of DSPE-PEG-Mal solution was added to the Tf solution. The reaction proceeded in PBS buffer (pH 7.0) for 2 h at room temperature under mild stirring. Similarly, GSH (Sigma Aldrich) and DSPE-PEG-MAL were incubated at 2:1 molar ration for 2 hours at RT (*34*). DSPE-PEG-Tf or DSPE-PEG-GSH were purified by dialysis.

### Preparation and characterization of NPs coated with different surface chemistries

siRNA-loaded PLGA NPs with different surface coatings were prepared by a modified nanoprecipitation method. Briefly, the organic phase was prepared by mixing 20 μL siRNA (4 nmol in water) with 1 mL PLGA (Durect Corporation) (5mg/mL in acetone, DMF, or tetrahydrofuran) and 200 μL of the cationic lipid-like molecule (5 mg/mL), which was synthesized by reacting the ethylenediamine core-poly (amidoamine) (PAMAM) generation 0 dendrimer (G0) (Sigma Aldrich) and 1,2 epoxytetradecane (Sigma Aldrich) using a ring opening reaction, as described previously (*56*). The siRNA sequences used in this study include: Luciferase siRNA: 5’-CUU ACG CUG AGU ACU UCG AdTdT-3’ (sense strand) and 5’-UCG AAG UAC UCA GCG UAA GdTdT-3’ (antisense strand); tau siRNA: 5’-CCU AGA AAU UCC AUG ACG AUU-3’ (sense strand) and 5’-UCG UCA UGG AAU UUC UAG GUU-3’ (antisense strand); scrambled siRNA: 5’-UUC UCC GAA CGU GUC ACG UUU-3’ (sense strand) and 5’-ACG UGA CAC GUU CGG AGA AUU-3’ (antisense strand); Cy3 or Dy677 labeled siRNAs were synthesized by labeling the 5’-end of both the sense and antisense strands of scrambled siRNA with Cy3 and Dy677. Under vigorous stirring, the mixture was added slowly into a 15 ml aqueous solution. Various coating materials were added to either the organic phase or water phase to endow the surface of NPs with different coatings. For PEG-NPs, the organic phase contained 2 mg/ml of DSPE-PEG (PEG molecular weight 3000) (Avanti Polar Lipids). For GSH-NPs, 5, 10 or 25 mol% of DSPE-PEG were replaced with DSPE-PEG-GSH to yield GSH (L)-NPs, GSH (M)-NPs, and GSH (H)-NPs, respectively. For Tf-NPs, 10 mol% of DSPE-PEG was replaced with DSPE-PEG-Tf. For F-68-NPs, the organic phase contained 1 mg/ml of DSPE-PEG, and the water phase contained 1 mg/ml Pluronic F-68 (Sigma Aldrich). For PS 80 (L)-NPs, the organic phase contained 1.5 mg/ml DSPE-PEG, and water phase contained 0.5 mg/ml PS 80 (Sigma Aldrich); for PS 80 (M)-NPs, the organic phase contained 1 mg/ml DSPE-PEG, and water phase contained 1 mg/ml PS 80; for PS 80 (H)-NPs, the water phase contained 2 mg/ml PS 80. Concentrations of surface coating materials were based on previously published reports (*34, 35*). PS 80-NPs and GSH-NPs used for evaluating the impact of surface coating chemistry on cellular uptake, gene silencing and BBB penetration were formulated with medium (M) coating densities of PS 80 and GSH. The resulting NP dispersions were transferred to centrifuge filters with 300kDa MWCO (Sartorius Vivaspin) and collected via centrifugation and washed with cold water three times. Finally, they were dispersed in PBS. Particle size distribution and surface charge of NPs were measured by dynamic light scattering (DLS, Brookhaven Instruments Corporation). NPs were stained with 1% uranyl acetate and their morphologies and size were observed under the Tecnai G2 Spirit BioTWIN transmission electron microscope (TEM, FEI Company).

To determine siRNA encapsulation efficiency, Cy3 labeled scrambled siRNA-loaded NPs were prepared as described above. 5 μL of NP solution was removed and mixed with 95 μL dimethyl sulfoxide (DMSO). Fluorescence intensity was analyzed using a Synergy HT multi-mode microplate reader (BioTek Instruments). Release kinetics of siRNA from NPs was also evaluated using Cy3 labeled scrambled siRNA-loaded NPs. Suspension of NPs in PBS was placed in dialysis tubing (float-a-lyzer G2 dialysis device, MWCO 100 kDa, SpectrumLabs) and dialyzed against PBS (pH 7.4) at 37 °C with a shaking speed of 150 rpm. At different time points, an aliquot of the NP suspension was withdrawn and dissolved in DMSO, and the fluorescence intensity of siRNA was analyzed. To determine PS 80 coating density on PS 80-coated NPs, PS 80-NP suspension was added to DMSO in a 1:10 ratio to dissolve the NPs and analyzed using high performance liquid chromatography-HPLC-evaporating light scattering detector (HPLC-ELSD) (Agilent 1260 Infinity II. Column: Zorbax 300SB-C18, 3 × 150 mm, 3.5 μm. Flow rate: 1 mL/min. PS 80 peak elution time = 7.8 min in a water-acetonitrile gradient). For GSH coating density calculation, we assumed a 100% incorporation of DSPE-PEG-GSH in GSH-NPs.

### Cell culture

Neuro-2a cells (ATCC® CCL-131™) were purchased from the American Type Culture Collection (ATCC). Cells were maintained in Eagle’s Minimum Essential Medium (EMEM) (ATCC) supplemented with 10% fetal bovine serum (Gibco) and 1% penicillin-streptomycin antibiotic (Thermo-Fisher Scientific). Luciferase-expressing Neuro-2a cells were generated by the transduction of cells with luciferase expression vector. The lentiviral vector pLenti CMV Puro LUC encoding firefly luciferase was transfected with Virapower Lentiviral packing mix to 293T cells (ATCC® CRL-3216™) using lipofectamine 2000 (Invitrogen). After 48 h, lentiviral supernatant was collected and added into 20-40% confluent Neuro-2a cells. Polybrene (8 μg/mL) was added during the transduction. Two days after the transduction, Neuro-2a cells were selected by puromycin (2 μg/mL) and were maintained in EMEM media containing 10% FBS and 1 μg/mL puromycin. All cells were incubated at 37 °C in a humidified atmosphere containing 5% CO_2_ in a cell culture incubator.

### Cellular uptake and endosomal escape of NPs

For cellular uptake study, Neuro-2a cells were seeded into the 8-well chambered coverglass (Nunc™ Lab-Tek™ Chambered Coverglass, Thermo Fisher Scientific) at the density of 15,000 cells/well. Cells were allowed to attach for 24 h. After that, cells were incubated with Dy677 labeled scrambled siRNA (free), or Dy677 labeled siRNA-loaded NPs having different surface coatings or coating densities. siRNA concentration was 15 nM. After incubation at 37 °C for 2 h, cells were washed twice with cold PBS buffer. Nuclei of cells were stained with Hoechst 33342 (Sigma Aldrich) (1 μg/mL) and cells imaged using CLSM (Olympus, FV1200). For endosomal escape, cells were incubated according to the procedures described above, except that before nuclear staining, cells were treated with green LysoTracker® Green (Life Technologies) at a final concentration of 60 nM for 30 min at 37 °C. Cells were then stained with Hoechst 33342 and observed under CLSM.

Uptake of NPs by Neuro-2a cells was quantified by flow cytometry. Neuro-2a cells were seeded in 6-well plates at 50,000 cells/well. After incubation at 37 °C for 2 days, cells were treated with Cy3 labeled scrambled siRNA (free) or Cy3 labeled siRNA-loaded NPs having different surface coatings or coating densities at final siRNA concentration of 15 nM. After 2 h, cells were washed twice with PBS buffer and then harvested by trypsin treatment for flow cytometry quantitative analysis (BD Biosciences FACS flow cytometer).

### In vitro gene silencing efficiency of NPs

Luciferase-expressing Neuro-2a cells were seeded in 96 well plates at 5,000 cells/well. Cells were incubated at 37 °C in a CO_2_ incubator overnight followed by treatment with luciferase siRNA-loaded NPs having different surface coatings and coating densities at final siRNA concentration of 0, 5, 10, and 25 nM. As controls, different concentrations of free luciferase siRNA or luciferase siRNA complexed with lipofectamine-2000 (Invitrogen) were used. Lipofectamine 2000-siRNA complex (siRNA-Lipo2K) was prepared according to the manufacturer’s suggested protocols. Medium was replaced after 24 h of incubation, and cells were continuously cultured in medium only for another 48 h. Cell viability was determined by quantifying cellular metabolic activity using alamarBlue assay (Thermo Fisher Scientific). Metabolic activity for each group was normalized to the metabolic activity of cells treated with medium only. Steady-Go luciferase assay kit (Promega) was used to measure the expression of luciferase in cells. Luciferase expression for each group was normalized to the luciferase expression of cells treated with medium only. Fluorescence intensity (for alamarBlue assay) and luminescence (for luciferase assay) of cells were recorded using a microplate reader.

### In vitro BBB penetration of NPs

bEnd.3 cells (ATCC) were grown on 1% gelatin-coated flasks at 37 °C, 5% CO_2_ in DMEM with 10% FBS and 1% penicillin/streptomycin. Then, the cells were seeded on 2% growth factor reduced Matrigel-coated 12 mm inserts with 0.4 μm pore polycarbonate membrane (Corning Transwell®, New York, NY) at a density of 80,000 cells/well. Medium was replaced every 2-3 days. Transendothelial electrical resistance (TEER) was measured using an EVOM resistance meter (World Precision Instruments, Sarasota, FL). After 1 week, penetration experiments were performed. Medium containing Dy677 labeled scrambled siRNA-loaded NPs with PEG or PS 80 (H) coating were added to the apical compartment. After 4 h, filter inserts were withdrawn from the receiver compartment. Aliquots from the basolateral compartments were collected and fluorescence was quantified using Infinite Pro 200 plate reader (Tecan). Empty filters without cells were used as control to determine the 100% penetration of NPs. Penetration of Dy677 labeled siRNA-loaded PS 80 (H)-NPs was also studied in the absence of serum in the cell culture medium. Anti-low density lipoprotein receptor-related protein 1 (LRP1) antibody (CST, Danvers, MA) was further used to block the lipoprotein receptor. Both compartments of the insert were incubated with 200 μM anti-LRP1 antibody in DMEM prior to NP introduction. After 1 h, NPs were introduced to the apical compartment. Penetration assay was then performed as described above.

### Animals

All *in vivo* experiments, except the isolation of primary neuronal cells were performed in 6-10 week-old male C57BL/6J mice (Jackson Laboratory). For isolation of primary neural cells, female, pregnant C57BL/6J mice at E18 were used as donors. Group sizes for each experimental design were determined based on the minimal number of animals needed to achieve a significant difference of *P*◻<◻0.05 between experimental groups for the primary outcome of the individual experiment using statistical analysis indicated in each figure legend. We randomly selected mice from the cage to assign them to different experimental groups. Experiments were performed in specific pathogen-free animal facilities at Boston Children’s Hospital (BCH). Mice were housed under standard 12◻h light–12◻h dark conditions with ad libitum access to water and chow. Animal care and handling procedures complied with the National Institutes of Health Guide for the Care and Use of Laboratory Animals. Animal protocols were approved by the Institutional Animal Care and Use Committee (IACUC) at BCH.

### Penetration of NPs across intact BBB in healthy mice

Dy677 labeled scrambled siRNA (free) or Dy677 labeled siRNA-loaded NPs having different surface coatings or coating densities were intravenously administrated into healthy male C57BL/6 mice *via* tail vein at siRNA dose of 50 nmol/kg. At 4 hours or 24 hours after the injection, mice were perfused with saline and euthanized. Brains were collected for *ex vivo* imaging using the IVIS spectrum imaging system (Xenogen Imaging Technologies) at the excitation and emission wavelengths of 675 nm and 740 nm, respectively.

### Pharmacokinetics of NPs

Healthy male C57BL/6 mice were intravenously injected with Dy677 labeled scrambled siRNA (free) or Dy677 siRNA-loaded PEG-NPs or PS 80 (H)-NPs *via* tail vein at siRNA dose of 50 nmol/kg. At pre-determined time points, blood was withdrawn from the tail artery and Dy677 siRNA fluorescence signal in blood was measured using a microplate reader (BioTek) at excitation and emission wavelengths of 670 and 710 nm, respectively.

### Isolation of primary neuronal cells and tau silencing in vitro

Primary neuronal cells were isolated from the cerebral cortex of mouse embryos. Briefly, pregnant female C57BL/6 mice at E18 were euthanized, the cortex was dissected from embryo brains, and meninges were removed thoroughly. Tissues were collected in a 15 mL conical tube and enzymatically digested at 37 °C for 20 min. The obtained cell solution was then passed through 40 μm cell strainer to remove tissue debris and centrifuged at 300 g for 4 min to collect the cells. Finally, cells were resuspended in B-27 Plus Neurobasal Medium (Thermo Fisher Scientific) and plated on culture plates pre-coated with poly-L-lysine (Sigma Aldrich). Half of the medium was changed every three days. After 10 days in culture, primary neuronal cells were incubated for 24 h with medium only (control) or medium containing free tau siRNA, tau siRNA-loaded PEG-NPs, scrambled (control) siRNA-loaded PS 80 (H)-NPs or tau siRNA-loaded PS 80 (H)-NPs. Following an additional 48 h incubation with fresh medium, cells were lysed with lysis buffer (Cell Signaling Technology) supplemented with protease inhibitor cocktail (Thermo Fisher Scientist) for 1 h on ice. Cell lysate was centrifuged at 14,000 rpm for 10 min, and supernatant was collected for measuring tau protein expression using western blot analysis. For western blotting, cell lysates from different treatment groups were resolved on SDS-PAGE gels (Thermo Fisher Scientific) and transferred to polyvinylidene difluoride (PVDF) membrane. Blots were blocked with 3% BSA in TBST (Tris Buffered Saline with Tween 20) for 1 h and then incubated with mouse anti-tau (Abcam) and rabbit anti-beta actin (Abcam) primary antibody at 4 °C overnight. After further incubation with horseradish peroxidase-conjugated goat anti-mouse IgG H&L and goat anti-rabbit IgG H&L (Thermo Fisher Scientific) for 1 hour, the protein bands were then visualized using chemiluminescent (ECL) detection reagent (Cell Signaling Technology) and imaged using a multi-application gel imaging system (Syngene). For quantification of tau expression, bands were analyzed using the Gels tool in Image J. The relative expression level of tau (tau/beta actin) was normalized to control group.

For immunofluorescence staining, primary neuronal cells were incubated with medium only (control) or medium containing free tau siRNA (10 nM siRNA), tau siRNA-loaded PEG-NPs (10 mM siRNA), or tau siRNA-loaded PS 80 (H)-NPs (5, 10 or 25 nM siRNA) for 24 h. Cells were further cultured in fresh medium for another 48 h, followed by fixation with 4% paraformaldehyde (PFA) for 10 min at room temperature. Cells were then permeabilized using 0.1% Triton X-100, and blocked by 1% BSA and 10% goat serum for 1 h. Cells were then incubated with mouse anti-tau primary antibody (10 μg/mL in 1% BSA) overnight at 4 °C, followed by incubation with Alexa Fluor® 488 goat anti-mouse IgG H&L (Abcam) (2 μg/mL) for 1 h at room temperature in dark. Nuclei were counterstained with Hoechst 33342. Finally, cells were imaged using CLSM (Olympus, FV1200).

### Traumatic brain injury

Mouse model of TBI was established using weight-drop method as reported previously (*27*). Male C57BL/6 mice were anesthetized using 4% isoflurane in oxygen and placed on a delicate task wiper (Kimwipe). Mice were then held by their tail and their heads were placed directly under a hollow guide tube. A 54 g metal bolt was dropped from a 60-inch height, which delivered an impact to the dorsal aspect of the skull and led to a rotational acceleration of head through KimWipe. All mice were allowed to recover in room air after injury and no mortality was found.

### Assessment of BBB permeability after TBI

BBB permeability of mice following weight drop induced TBI was assessed by EB penetration assay. EB dye (2% w/v) was intravenously injected into healthy mice or mice with TBI at a dose of 4 ml/kg. Mice with TBI received EB at different time points (5 min, 6 h, 24 h, 1 week, and 2 weeks) post-injury. Two hours after EB administration, animals were perfused extensively with saline and then euthanized to collect the whole brain. Photos of brains were taken with a digital camera. To quantify the amount of EB in the brain, brain tissue was cut into small pieces and homogenized. EB was extracted from brain homogenate by treatment with 60% trichloroacetic acid (TCA). Fluorescence intensity of EB was recorded using a plate reader at an excitation wavelength of 620 nm and an emission wavelength of 680 nm.

### Penetration of NPs across BBB in TBI mice

Dy677 labeled scrambled siRNA (free) or Dy677 labeled siRNA-loaded PEG-NPs or PS 80 (H)-NPs were administered intravenously into TBI mice at siRNA dose of 50 nmol/kg, either at 2 h or 2 weeks after injury. At 4 h post-injection, mice were anesthetized, and perfused with saline. Brains were harvested and imaged with IVIS Spectrum imaging system (Xenogen). To specifically localize delivered siRNA within brain tissue, blood vessels were labeled with FITC-lectin (Vector Laboratories) by injecting it into mice *via* tail vein 10 min before euthanasia (*57*). Mice were then anesthetized, and perfused with saline and 4% PFA. Next, brains were harvested, and fixed in 4% PFA. Brains were further equilibrated in 30% sucrose solution and transferred to tissue base mold. After covering the tissue block with Optimum Cutting Temperature Compound (OCT, Fisher Scientist), base molds containing tissue blocks were immersed into liquid nitrogen until the tissue was frozen completely. Frozen brain tissues were cut into 40 μm sections and counterstained with Hoechst 33342, followed by imaging with a fluorescent microscope.

### Tau silencing in vivo

TBI inducing weight drop procedure was performed on day 0. To evaluate tau silencing efficiency during early injury, mice received tail vein injection of PBS, free tau siRNA, tau siRNA-loaded PS 80 (H) NPs, or scrambled (control) siRNA-loaded PS 80 (H)-NPs at siRNA dose of 75 nmol/kg/day at 2 h and 1 day post-injury. To evaluate tau silencing efficiency during late injury, a separate group of mice received tail vein injections on day 14 and 15 post-injury. Three days after the last injection in each injury period, mice were sacrificed, and brains were harvested to determine tau expression in cortex region using western blot. For 10 mg cortex tissue, 500 μL of ice-cold lysis buffer was added. Tissues were homogenized and the homogenate was centrifuged for 20 min at 14,000 rpm at 4 °C. The supernatant was carefully collected and analyzed by western blotting as described above. The relative expression level of tau (tau/beta actin) was normalized to naïve animal group (healthy mice with no treatment).

For immunohistochemistry staining, harvested brain tissues were subjected to PFA-fixed paraffin-embedded section. The paraffin-embedded sections (5-μm thick) were deparaffinized with xylene, and rehydrated by immersing the slides in a series of different concentrations of alcohol. Following antigen retrieval using DAKO target retrieval solution, samples were incubated with DAKO peroxidase blocking buffer at room temperature for 10 min to quench endogenous peroxidase activity. Slides were then incubated with the anti-tau primary antibody solution, followed by incubation with peroxidase-labeled polymer conjugated to secondary antibodies for 30 min. Slides were finally stained with DAB+ substrate-chromogen solution and hematoxylin and imaged using Aperio digital slide scanner (20X).

### In vivo safety

Healthy C57BL/6 mice were intravenously injected with PBS (control) or PS 80 (H)-NPs loaded with scrambled (control) siRNA or tau siRNA at siRNA dose of 75 nmol/kg/day for 2 consecutive days. Main organs including lung, heart, liver, spleen, and kidney were collected 3 days after the last dose and fixed in 4% PFA, followed by embedding in paraffin and sectioning for hematoxylin and eosin (H&E) staining. H&E sections (5 μm) were imaged using the Aperio digital slide scanner (20X). In addition to H&E staining, the hematology markers and multiple biochemical parameters in blood were also evaluated. Specifically, 3 days or 2 weeks after the last dose, mice were euthanized, and blood was drawn by a retro-orbital puncture. 1 mL blood was collected in the Microvette® blood collection tube and analyzed within 1 h using automated hematology analyzer. Hematological parameters examined in this study included white blood cells (WBC), red blood cells (RBC), hemoglobin (HGB), hematocrit (HCT), mean corpuscular hemoglobin (MCH), platelets (PLT), mean platelet volume (MPV), red cell distribution width (RDW), mean corpuscular hemoglobin concentration (CHC), mean corpuscular volume (MCV). For biochemical analysis, serum was collected by Microvette® serum gel tube and analyzed for alkaline phosphatase (ALP), alanine aminotransferase (ALT), aspartate aminotransferase (AST), creatinine, and blood urea nitrogen (BUN).

### Statistics

Statistical analysis and graphing were done with Graphpad Prism. The two-tailed Student’s t-test was used to compare two experimental groups and one-way ANOVA with Tukey’s post hoc analysis was used for comparing more than two groups. A value of *P* < 0.05 was considered statistically significant.

## Acknowledgments

This work was supported by the National Institutes of Health grant HL095722 (to J.M.K.) and Fundação para a Ciência e a Tecnologia through MIT-Portugal-TB/ECE/0013/2013 (to J.M.K.). This work was also conducted with the support of the Football Players Health Study at Harvard (to R.M). The Football Players Health Study is funded by a grant from the National Football League Players Association. The content is solely the responsibility of the authors and does not necessarily represent the official views of Harvard Medical School, Harvard University or its affiliated academic health care centers, the National Football League Players Association, or the Brigham and Women’s Hospital.

## Author contributions

W.L, J.Q., R.M., J.M.K. and N.J. conceived the study and designed experiments. W.L., J.Q., X.L., S.A., J.Z., G.C. and J.X. performed the experiments and analyzed the data with N.J. and J.M.K. W.L. crafted all the figures and wrote the manuscript. X.L. performed all the statistical analysis. R.L., R.M., J.M.K. and N.J. edited and revised the manuscript and supervised the research.

## Competing interests

JMK has been a paid consultant and or equity holder for multiple biotechnology companies (listed here https://www.karplab.net/team/jeff-karp). The interests of JMK were reviewed and are subject to a management plan overseen by his institutions in accordance with its conflict of interest policies. N.J., J.MK., W.L., R.M., J.Q. and R.L. have one unpublished patent based on the nanoparticle work presented in this manuscript. For a list of entities with which R.L. is involved, compensated or uncompensated, see www.dropbox.com/s/yc3xqb5s8s94v7x/Rev%20Langer%20COI.pdf?dl=0.

## Data and materials availability

All data associated with this study are present in the paper or in the Supplementary Materials.

## References and Notes

1. B. Roozenbeek, A. I. R. Maas, D. K. Menon, Changing patterns in the epidemiology of traumatic brain injury. Nat. Rev. Neurol. 9, 231–236 (2013).

2. D. W. Simon, M. J. McGeachy, H. Bayır, R. S. B. Clark, D. J. Loane, P. M. Kochanek, The far-reaching scope of neuroinflammation after traumatic brain injury. Nat. Rev. Neurol. 13, 171–191 (2017).

3. J. S. Meabon, B. R. Huber, D. J. Cross, T. L. Richards, S. Minoshima, K. F. Pagulayan, G. Li, K. D. Meeker, B. C. Kraemer, E. C. Petrie, M. A. Raskind, E. R. Peskind, D. G. Cook, Repetitive blast exposure in mice and combat veterans causes persistent cerebellar dysfunction. Sci. Transl. Med. 8, 321ra6 (2016)

4. L. E. Goldstein, A. M. Fisher, C. A. Tagge, X.-L. Zhang, L. Velisek, J. A. Sullivan, C. Upreti, J. M. Kracht, M. Ericsson, M. W. Wojnarowicz, C. J. Goletiani, G. M. Maglakelidze, N. Casey, J. A. Moncaster, O. Minaeva, R. D. Moir, C. J. Nowinski, Robert A. Stern, R. C. Cantu, J. Geiling, J. K. Blusztajn, B. L. Wolozin, T. Ikezu, T. D. Stein, A. E. Budson, N. W. Kowall, D. Chargin, A. Sharon, S. Saman, G. F. Hall, W. C. Moss, R. O. Cleveland, R. E. Tanzi, P. K. Stanton, A. C. McKee, Chronic traumatic encephalopathy in blast-exposed military veterans and a blast neurotrauma mouse model. Sci. Transl. Med. 4, 134ra160 (2012).

5. A. C. McKee, D. Stein, C. J. Nowinski, R. A. Stern, D. H. Daneshvar, V. E. Alvarez, H.-S. Lee, G. Hall, S. M. Wojtowicz, C. M. Baugh, D. O. Riley, C. A. Kubilus, K. A. Cormier, M. A. Jacobs, B. R. Martin, C. R. Abraham, T. Ikezu, R. R. Reichard, B. L. Wolozin, A. E. Budson, L. E. Goldstein, N. W. Kowall, R. C. Cantu, The spectrum of disease in chronic traumatic encephalopathy. Brain 136, 43–64 (2012).

6. D. H. Smith, V. E. Johnson, W. Stewart, Chronic neuropathologies of single and repetitive TBI: substrates of dementia? Nat. Rev. Neurol. 9, 211–221 (2013).

7. S. T. DeKosky, K. Blennow, M. D. Ikonomovic, S. Gandy, Acute and chronic traumatic encephalopathies: pathogenesis and biomarkers. Nat. Rev. Neurol. 9, 192–200 (2013).

8. K. Blennow, J. Hardy, H. Zetterberg, The neuropathology and neurobiology of traumatic brain injury. Neuron 76, 886–899 (2012).

9. D. J. Sharp, G. Scott, R. Leech, Network dysfunction after traumatic brain injury. Nat. Rev. Neurol. 10, 156–166 (2014).

10. D. Yoo, A. W. Magsam, A. M. Kelly, P. S. Stayton, F. M. Kievit, A. J. Convertine, Core-cross-linked nanoparticles reduce neuroinflammation and improve outcome in a mouse model of traumatic brain injury. ACS Nano, 11, 8600 (2017).

11. K. A. Whitehead, R. Langer, D. G. Anderson, Knocking down barriers: advances in siRNA delivery. Nat. Rev. Drug Discov. 8, 129–138 (2009).

12. S. F. Dowdy, Overcoming cellular barriers for RNA therapeutics. Nat. Biotechnol. 35, 222–229 (2017).

13. M. Zheng, W. Tao, Y. Zou, O. C. Farokhzad, B. Shi, Nanotechnology-based strategies for siRNA brain delivery for disease therapy. Trends Biotechnol. 36, 562–575 (2018).

14. E. J. Kwon, M. Skalak, R. L. Bu, S. N. Bhatia, Neuron-targeted nanoparticle for siRNA delivery to traumatic brain injuries. ACS Nano 10, 7926–7933 (2016).

15. J. Kang, J. Joo, E. J. Kwon, M. Skalak, S. Hussain, Z.-G. She, E. Ruoslahti, S. N. Bhatia, M. J. Sailor, Self-sealing porous silicon-calcium silicate core–shell nanoparticles for targeted siRNA delivery to the injured brain. Adv. Mater. 28, 7962–7969 (2016).

16. V. N. Bharadwaj, J. Lifshitz, P. D. Adelson, V. D. Kodibagkar, S. E. Stabenfeldt, Temporal assessment of nanoparticle accumulation after experimental brain injury: Effect of particle size. Sci. Rep. 6, 29988 (2016).

17. Y. Xiong, A. Mahmood, M. Chopp, Animal models of traumatic brain injury. Nat. Rev. Neurosci. 14, 128–142 (2013).

18. A. Chodobski, B. J. Zink, J. Szmydynger-Chodobska, Blood-brain barrier pathophysiology in traumatic brain injury. Transl. Stroke Res. 2, 492–516 (2011).

19. M. Kuriakose, K. V. R. Rao, D. Younger, N. Chandra, Temporal and spatial effects of blast overpressure on blood-brain barrier permeability in traumatic brain injury. Sci. Rep. 8, 8681 (2018).

20. R. Sahyouni, P. Gutierrez, E. Gold, R. T. Robertson, B. J. Cummings, Effects of concussion on the blood– brain barrier in humans and rodents. J. Concussion. 1, 1–15 (2017).

21. M. Nowak, M. E. Helgeson, S. Mitragotri, Delivery of nanoparticles and macromolecules across the Blood Brain Barrier. Adv. Therap., 3, 1900073 (2019)

22. D. Furtado, M. Björnmalm, S. Ayton, A. I. Bush, K. Kempe, F. Caruso, Overcoming the blood–brain barrier: the role of nanomaterials in treating neurological diseases. Adv. Mater. 30, 1801362 (2018).

23. Y. Anraku, H. Kuwahara, Y. Fukusato, A. Mizoguchi, T. Ishii, K. Nitta, Y. Matsumoto, K. Toh, K. Miyata, S. Uchida, K. Nishina, K. Osada, K. Itaka, N. Nishiyama, H. Mizusawa, T. Yamasoba, T. Yokota, K. Kataoka Glycaemic control boosts glucosylated nanocarrier crossing the BBB into the brain. Nat. Commun. 8, 1001 (2017).

24. J.-L. Huang, G. Jiang, Q.-X. Song, X. Gu, M. Hu, X.-L. Wang, H.-H. Song, L.-P. Chen, Y.-Y. Lin, D. Jiang, J. Chen, J.-F. Feng, Y.-M. Qiu, J.-Y. Jiang, X.-G. Jiang, H.-Z. Chen, X.-L. Gao, Lipoprotein-biomimetic nanostructure enables efficient targeting delivery of siRNA to Ras-activated glioblastoma cells via macropinocytosis. Nat. Commun. 8, 15144 (2017).

25. J. Hrkach, D. Von Hoff, M. M. Ali, E. Andrianova, J. Auer, T. Campbell, D. De Witt, M. Figa, M. Figueiredo, A. Horhota, S. Low, K. McDonnell, E. Peeke, B. Retnarajan, A. Sabnis, E. Schnipper, J. J. Song, Y. H. Song, J. Summa, D. Tompsett, G. Troiano, T. Van Geen Hoven, J. Wright, P. LoRusso, P. W. Kantoff, N. H. Bander, C. Sweeney, O. C. Farokhzad, R. Langer, S. Zale, Preclinical development and clinical translation of a PSMA-targeted docetaxel nanoparticle with a differenciated pharmacological profile. Sci. Transl. Med. 4, 128ra139 (2012).

26. K. J. McHugh, T. D. Nguyen, A. R. Linehan, D. Yang, A. M. Behrens, S. Rose, Z. L. Tochka, S. Y. Tzeng, J. J. Norman, A. C. Anselmo, X. Xu, S. Tomasic, M. A. Taylor, J. Lu, R. Guarecuco, R. S. Langer, A. Jaklenec, Fabrication of fillable microparticles and other complex 3D microstructures. Science 357, 1138–1142 (2017).

27. A. Kondo, K. Shahpasand, R. Mannix, J. Qiu, J. Moncaster, C.-H. Chen, Y. Yao, Y.-M. Lin, J. A. Driver, Y. Sun, S. Wei, M.-L. Luo, O. Albayram, P. Huang, A. Rotenberg, A. Ryo, L. E. Goldstein, A. Pascual-Leone, A. C. McKee, W. Meehan, X. Z. Zhou, K. P. Lu, Antibody against early driver of neurodegeneration cis P-tau blocks brain injury and tauopathy. Nature 523, 431–436 (2015).

28. R. Mannix, W. P. Meehan, J. Mandeville, P. E. Grant, T. Gray, J. Berglass, J. Zhang, J. Bryant, S. Rezaie, J. Y. Chung, N. V. Peters, C. Lee, L. W. Tien, D. L. Kaplan, M. Feany, M. Whalen, Clinical correlates in an experimental model of repetitive mild brain injury. Ann Neurol. 74, 65–75 (2013).

29. J. S. Suk, Q. Xu, N. Kim, J. Hanes, L. M. Ensign, PEGylation as a strategy for improving nanoparticle-based drug and gene delivery. Adv. Drug Deliv. Rev. 99, 28–51 (2016).

30. J. Li, P. Cai, A. Shalviri, J. T. Henderson, C. He, W. D. Foltz, P. Prasad, P. M. Brodersen, Y. Chen, R. DaCosta, A. M. Rauth, X. Y. Wu, A multifunctional polymeric nanotheranostic system delivers doxorubicin and imaging agents across the blood–brain barrier targeting brain metastases of breast cancer. ACS Nano 8, 9925–9940 (2014).

31. J. Li, P. Cai, A. Shalviri, J. T. Henderson, C. He, W. D. Foltz, P. Prasad, P. M. Brodersen, Y. Chen, R. DaCosta, A. M. Rauth, X. Y. Wu, Nanoparticles enhance brain delivery of blood–brain barrier-impermeable probes for in vivo optical and magnetic resonance imaging. Proc. Natl. Acad. Sci. U.S.A. 108, 18837–18842 (2011).

32. M. Naito, T. Yokoyama, K. Hosokawa, K. Nogi, Nanoparticle technology handbook 3rd edition, 471 (Elsevier, 2018).

33. D. Maussang, J. Rip, J. van Kregten, A. van den Heuvel, S. van der Pol, B. van der Boom, A. Reijerkerk, L. Chen, M. de Boer, P. Gaillard, H. de Vries Glutathione conjugation dose-dependently increases brain-specific liposomal drug delivery in vitro and in vivo. Drug Discov. Today: Technol. 20, 59–69 (2016).

34. J. Rip, L. Chen, R. Hartman, A. van den Heuvel, A. Reijerkerk, J. van Kregten, B. van der Boom, C. Appeldoorn, M. de Boer, D. Maussang, E. C. M. de Lange, P. J. Gaillard, Glutathione PEGylated liposomes: pharmacokinetics and delivery of cargo across the blood–brain barrier in rats. J. Drug Target. 22, 460–467 (2014).

35. F. C. Lam, S. W. Morton, J. Wyckoff, T.-L. V. Han, M. K. Hwang, A. Maffa, E. Balkanska-Sinclair, M. B. Yaffe, S. R. Floyd, P. T. Hammond, Enhanced efficacy of combined temozolomide and bromodomain inhibitor therapy for gliomas using targeted nanoparticles. Nat. Commun. 9, 1991 (2018).

36. B. Oller-Salvia, M. Sánchez-Navarro, E. Giralt, M. Teixidó, Blood–brain barrier shuttle peptides: an emerging paradigm for brain delivery. Chem. Soc. Rev. 45, 4690–4707 (2016).

37. C. Saraiva, C. Praça, R. Ferreira, T. Santos, L. Ferreira, L. Bernardino, Nanoparticle-mediated brain drug delivery: Overcoming blood–brain barrier to treat neurodegenerative diseases. J. Control. Release 235, 34–47 (2016).

38. N. Hoshyar, S. Gray, H. Han, G. Bao, The effect of nanoparticle size on in vivo pharmacokinetics and cellular interaction. Nanomedicine 11, 673–692 (2016).

39. G. Sahay, W. Querbes, C. Alabi, A. Eltoukhy, S. Sarkar, C. Zurenko, E. Karagiannis, K. Love, D. Chen, R. Zoncu, Y. Buganim, A. Schroeder, R. Langer, D. G. Anderson, Efficiency of siRNA delivery by lipid nanoparticles is limited by endocytic recycling. Nat. Biotechnol. 31, 653 (2013).

40. J. P. J. Wu, B. Cheng, S. R. Roffler, D. J. Lundy, C. Y. T. Yen, P. Chen, J. J. Lai, S. H. Pun, P. S. Stayton, P. C. H. Hsieh, Reloadable multidrug capturing delivery system for targeted ischemic disease treatment. Sci. Transl. Med., 8, 365ra160 (2016)

41. E. A. Nance, G. F. Woodworth, K. A. Sailor, T. Y. Shih, Q. Xu, G. Swaminathan, D. Xiang, C. Eberhart, J. Hanes, A dense poly(ethylene glycol) coating improves penetration of large polymeric nanoparticles within brain tissue. Sci. Transl. Med. 4, 149ra119 (2012)

42. A. Ambruosi, A. S. Khalansky, H. Yamamoto, S. E. Gelperina, D. J. Begley, J. Kreuter, Biodistribution of polysorbate 80-coated doxorubicin-loaded [14C]-poly(butyl cyanoacrylate) nanoparticles after intravenous administration to glioblastoma-bearing rats. J. Drug Target. 14, 97–105 (2006).

43. H. Jaffer, I. M. Adjei, V. Labhasetwar, Optical imaging to map blood-brain barrier leakage. Sci. Rep. 3, 3117 (2013).

44. P. Shahim, Y. Tegner, D. Wilson, J. Randall, B. Kallberg, K. Blennow, H. Zetterberg, Blood biomarkers for brain injury in concussed professional ice hockey players. JAMA Neurol. 71, 684–692 (2014).

45. J. S. Cheng, R. Craft, G.-Q. Yu, K. Ho, X. Wang, G. Mohan, S. Mangnitsky, R. Ponnusamy, L. Mucke, Tau reduction diminishes spatial learning and memory deficits after mild repetitive traumatic brain injury in mice. PLoS ONE 9, e115765 (2014).

46. S. L. DeVos, R. L. Miller, K. M. Schoch, B. B. Holmes, C. S. Kebodeaux, A. J. Wegener, G. Chen, T. Shen, H. Tran, B. Nichols, T. A. Zanardi, H. B. Kordasiewicz, E. E. Swayze, C. F. Bennett, M. I. Diamond, T. M. Miller, Tau reduction prevents neuronal loss and reverses pathological tau deposition and seeding in mice with tauopathy. Sci. Transl. Med. 9, eaag0481 (2017).

47. H. Xu, T. W. Rösler, T. Carlsson, A. de Andrade, O. Fiala, M. Höllerhage, W. H. Oertel, M. Goedert, A. Aigner, G. U. Höglinger, Tau silencing by siRNA in the P301S mouse model of tauopathy. Curr. Gene Ther. 14, 343–51 (2014).

48. M. Chiasseu, J. L. C. Vargas, L. Destroismaisons, C. V. Velde, N. Leclerc, A. D. Polo, Tau accumulation, altered phosphorylation, and missorting promote neurodegeneration in glaucoma. J. Neurosci. 36, 5785–5798 (2016).

49. S. F. Carron, D. S. Alwis, R. Rajan, Traumatic brain injury and neuronal functionality changes in sensory cortex. Front. Syst. Neurosci. 10, 47 (2016).

50. A. M. Fukuda, J. Badaut, siRNA treatment: “a sword-in-the-stone” for acute brain injuries. Genes 4, 435–456 (2013).

51. N. Marklund, L. Hillered, Animal modelling of traumatic brain injury in preclinical drug development: where do we go from here? Br. J. Pharmacol. 164, 1207–1229 (2011).

52. L. Calcul, B. Zhang, U. K. Jinwal, C. A. Dickey, B. J. Baker, Natural products as a rich source of tau-targeting drugs for Alzheimer’s disease. Future Med. Chem. 4, 1751–1761 (2012).

53. A. M. P. Siahaan, I. Japardi, A. S. Rambe, R. S. Indharty, M. Ichwan, Turmeric extract supplementation reduces Tau protein level in repetitive traumatic brain injury model. J. Med. Sci. 6, 1953–1958 (2018).

54. C. Zheng, C. Ma, E. Bai, K. Yang, R. Xu, Transferrin and cell-penetrating peptide dual-functioned liposome for targeted drug delivery to glioma. Int. J. Clin. Exp. Med. 8, 1658–1668 (2015).

55. H. Liu, K. D. Moynihan, Y. Zheng, G. L. Szeto, A. V. Li, B. Huang, D. S. V. Egeren, C. Park, D. J. Irvine, Structure-based programming of lymph-node targeting in molecular vaccines. Nature 507, 519–522 (2014).

56. X. Xu, K. Xie, X.-Q. Zhang, E. M. Pridgen, G. Y. Park, D. S. Cui, J. Shi, J. Wu, P. W. Kantoff, S. J. Lippard, R. Langer, G. C. Walker, O. C. Farokhzad, Enhancing tumor cell response to chemotherapy through nanoparticle-mediated codelivery of siRNA and cisplatin prodrug. Proc. Natl. Acad. Sci. USA. 110, 18638–18643 (2013).

57. R. T. Robertson, S. T. Levine, S. M. Haynes, P. Gutierrez, J. L. Baratta, Z. Tan, K. J. Longmuir, Use of labeled tomato lectin for imaging vasculature structures. Histochem. Cell Biol. 143, 225–34 (2015).

