## Supplemental figures and tables for "BBB pathophysiology independent delivery of siRNA in traumatic brain injury"

Wen Li^1,2,3^, Jianhua Qiu^3,4^, Xiang-Ling Li^1^, Sezin Aday^1,2^, Jingdong Zhang^4,5^, Grace Conley^4^, Jun Xu^1,3^, Robert Langer^2,6^, Rebekah Mannix^3,4,*^, Jeffrey M. Karp^1,3,6,7,8,*^, Nitin Joshi^1,2,3,*^

^1^Center for Nanomedicine and Division of Engineering in Medicine, Department of Medicine, Brigham and Women’s Hospital, Boston, MA 02115, USA.

^2^Department of Chemical Engineering and Koch Institute for Integrative Cancer Research, Massachusetts Institute of Technology, Cambridge, MA 02139, USA.

^3^Harvard Medical School, Boston, MA 02115, USA.

^4^Division of Emergency Medicine, Boston Children’s Hospital, Boston, MA 02115, USA.

^5^Department of Neurosurgery, Brigham and Women’s Hospital, Boston MA 02115 USA.

^6^Harvard–Massachusetts Institute of Technology Division of Health Sciences and Technology, Massachusetts Institute of Technology, Cambridge, MA 02139, USA

^7^Broad Institute, Cambridge, MA 02142, USA

^8^Harvard Stem Cell Institute, Cambridge, MA 02138, USA

**Supplementary Figures**

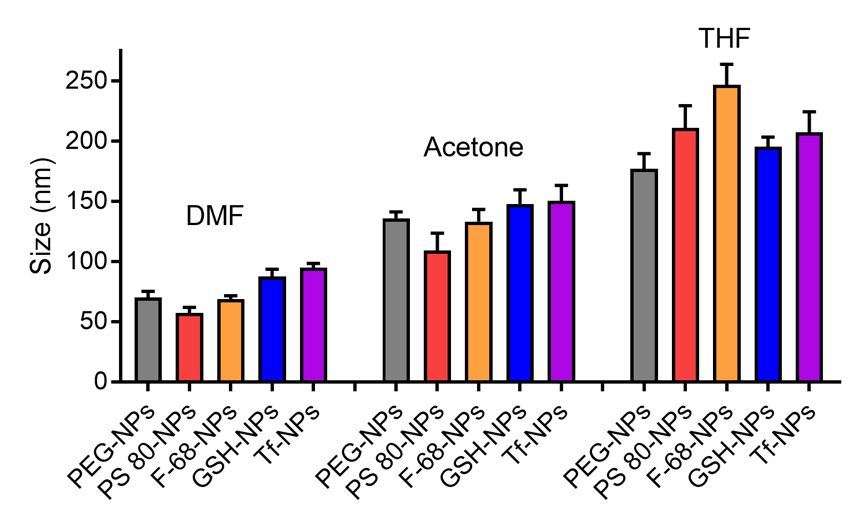

**Fig. S1. Size of siRNA-loaded NPs with different surface coatings analyzed by DLS.** Different organic solvents, including *N,N'*-dimethylformamide (DMF), acetone, or tetrahydrofuran (THF) were used for preparing the siRNA-loaded NPs. Data are mean ± SD of technical repeats (*n* = 3, experiment performed at least twice).

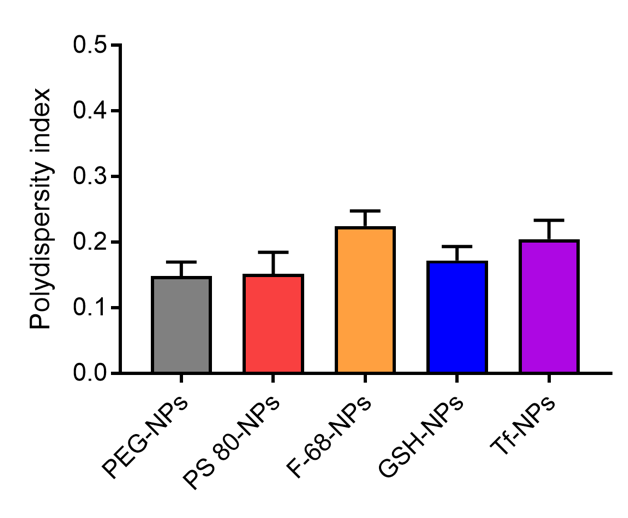

**Fig. S2.** **Polydispersity index of siRNA-loaded NPs with different surface coatings.** Polydispersity index of siRNA-loaded NPs was measured by DLS analysis. Data are mean ± SD of technical repeats (*n* = 3, experiment performed at least twice).

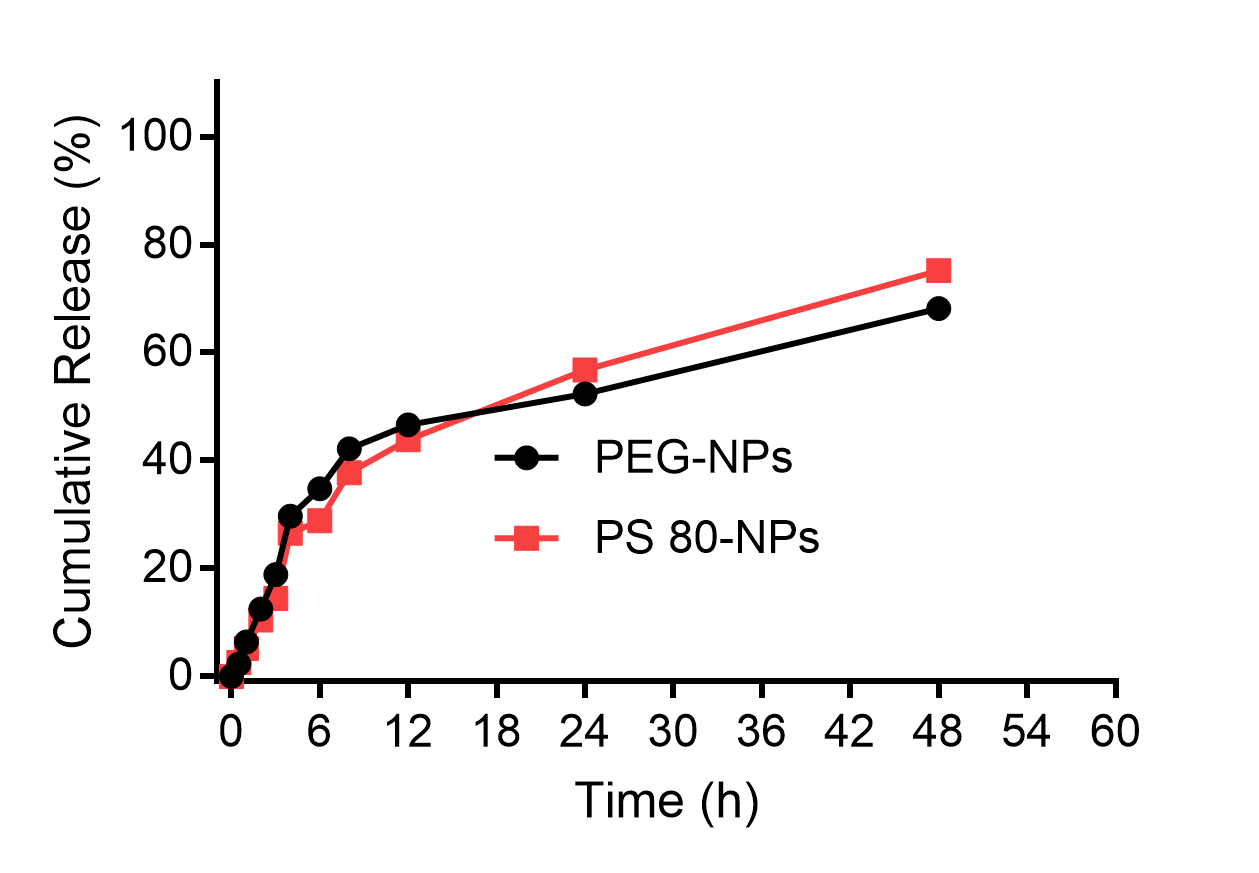

**Fig. S3.** ***In vitro* release profiles of siRNA from PEG-NPs or PS 80-NPs in PBS at 37°C.** Cy3 labelled scrambled siRNA was used for these experiments. Data are mean ± SD of technical repeats (*n* = 3, experiment performed at least twice).

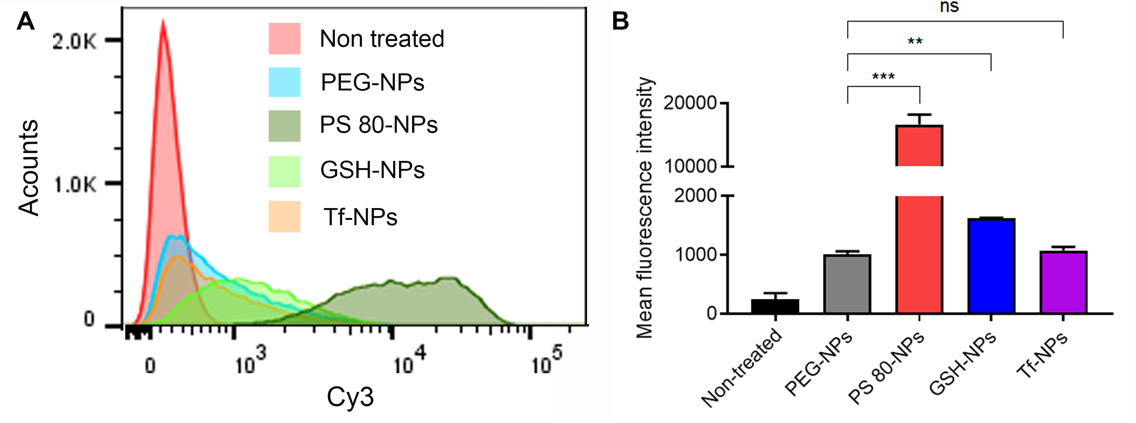

**Fig. S4.** **Cellular uptake of siRNA-loaded NPs having different surface coatings analyzed by flow cytometry.** (**A**) Flow cytometry histogram of Neuro-2a cells incubated at 37 °C for 2 h with medium only (non-treated) or medium containing siRNA-loaded NPs having different coatings (PEG-NPs, PS 80-NPs, GSH-NPs or Tf-NPs). Cy3 labelled scrambled siRNA was used. (**B**) Mean fluorescence intensity data from flow cytometry. ***P* < 0.01, *** *P* < 0.001. Data in B are mean ± SD of technical repeats (*n* = 3, experiment performed at least twice). *P*-values were determined using two-tailed Student’s *t* test.

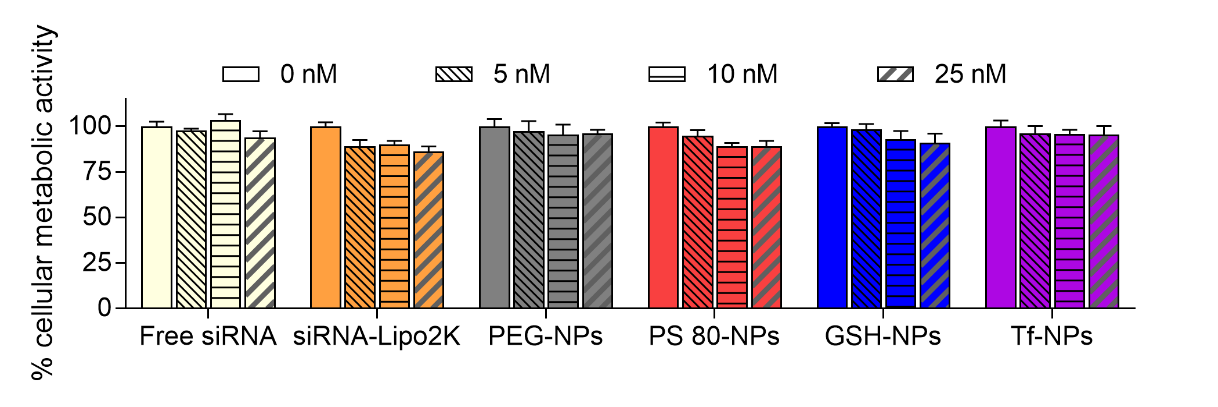

**Fig. S5**. **Viability of Neuro-2a cells after incubation with luciferase siRNA formulations.** Cell viability was determined by quantifying cellular metabolic activity using alamarBlue assay. Neuro-2a cells were incubated for 24 h with medium only or medium containing free luciferase siRNA, lipofectamine 2000-luciferase siRNA complex (siRNA-Lipo2K) or luciferase siRNA-loaded NPs having different surface coatings (PEG-NPs, PS 80-NPs, GSH-NPs or Tf-NPs). siRNA dose was 0, 5, 10, or 25 nM. Following an additional 48 h incubation with medium only, metabolic activity was determined using alamarBlue assay and normalized to the metabolic activity of cells treated with medium only (0 nM siRNA). Data are mean ± SD of technical repeats (*n* = 3, experiment performed at least twice).

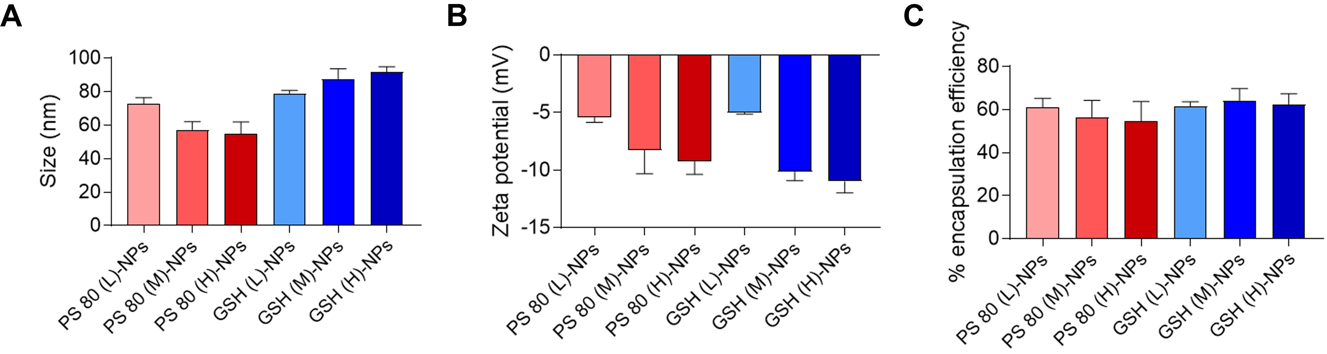

**Fig. S6.** **Characterization of siRNA-loaded NPs with different coating densities of PS 80 or GSH.** (**A**) Size and (**B**) zeta potential of siRNA-loaded NPs with different coating densities of PS 80 or GSH. (**C**) Encapsulation efficiency of Cy3 labelled scrambled siRNA in NPs with different coating densities of PS 80 or GSH. Data in A-C are mean ± SD of technical repeats (*n* = 3, experiment performed at least twice).

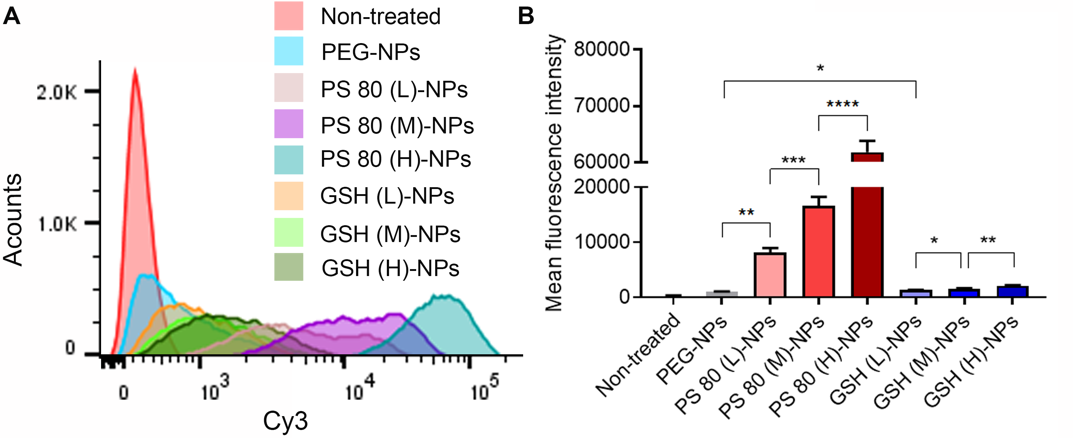

**Fig. S7.** **Cellular uptake of siRNA-loaded NPs having different coating densities of PS 80 or GSH analyzed by flow cytometry.** (**A**) Flow cytometry analysis of Neuro-2a cells incubated at 37 °C for 2 h with medium only (non-treated) or medium containing siRNA-loaded PEG-NPs or siRNA-loaded NPs having different coating densities of PS 80 or GSH. Cy3 labelled scrambled siRNA was used. (**B**) Mean fluorescence intensity data from flow cytometry. **P* < 0.05, ***P* < 0.01, ****P* < 0.001, *****P* < 0.0001. Data in B are mean ± SD of technical repeats (*n* = 3, experiment performed at least twice). *P*-values were determined using one-way ANOVA with Tukey’s post-hoc analysis.

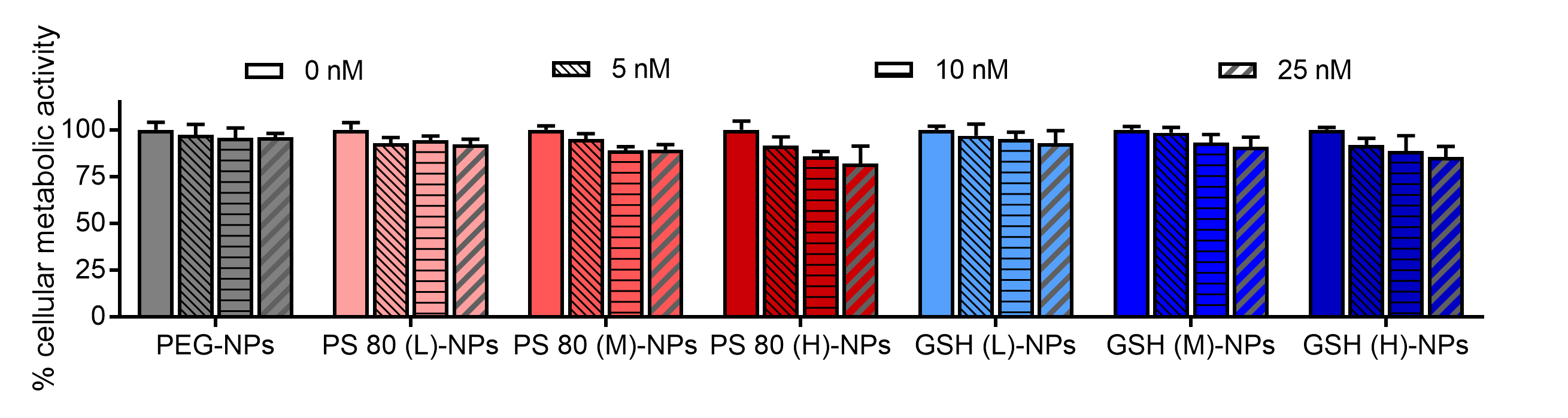

**Fig. S8. Viability of Neuro-2a cells after incubation with luciferase siRNA formulations.** Cell viability was determined by quantifying cellular metabolic activity using alamarBlue assay. Neuro-2a cells were incubated for 24 h with medium only or medium containing luciferase siRNA-loaded PEG NPs, or luciferase siRNA-loaded NPs having different coating densities of PS 80 or GSH. siRNA dose was 0, 5, 10, or 25 nM. Following an additional 48 h incubation with medium only, metabolic activity was determined using alamarBlue assay and normalized to the metabolic activity of cells treated with medium only (0 nM siRNA). Data in are mean ± SD of technical repeats (*n* = 3, experiment performed at least twice).

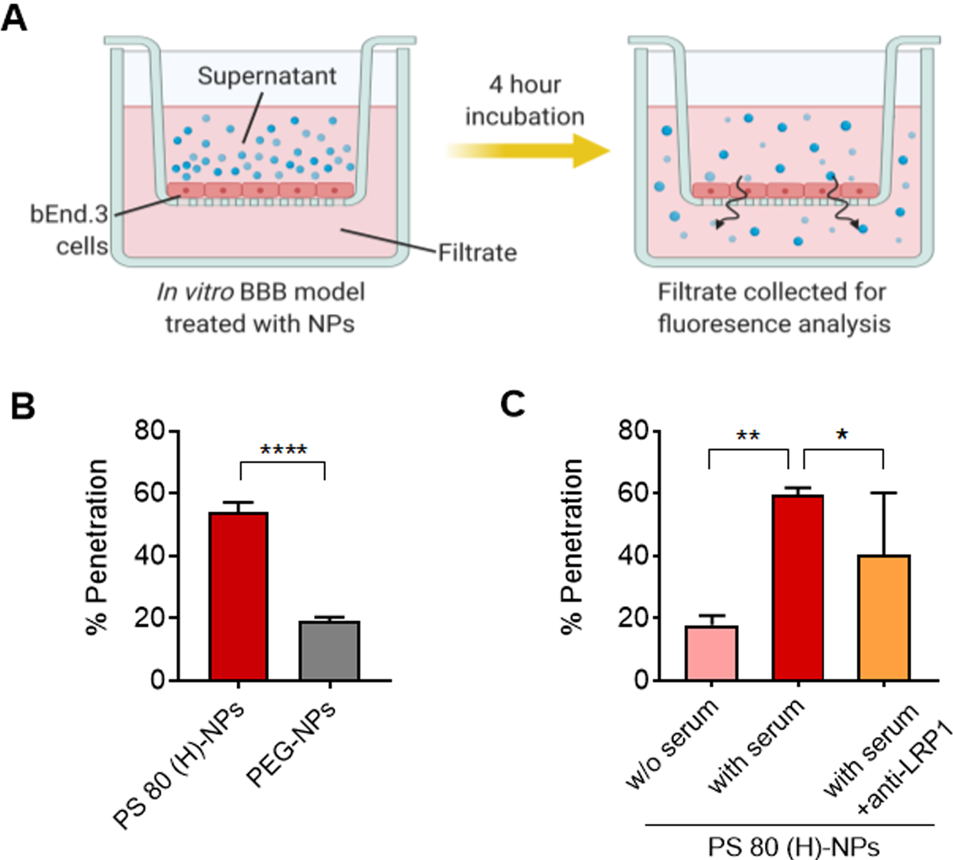

**Fig. S9.** **Evaluation of BBB-penetration ability of NPs in *in vitro* BBB model.** (**A**) Schematic depicting the assay used to determine the penetration of NPs through an *in vitro* BBB model. Mouse bEnd.3 cell monolayers were incubated for 4 h with Dy677 labeled scrambled siRNA-loaded PEG-NPs or PS 80 (H)-NPs, added to the apical side. Fluorescence in the filtrate was then measured to determine the fraction of NPs that had penetrated the monolayer. (**B**) Penetration of NPs through the cell monolayer was calculated by normalizing the fraction of NPs that penetrated the monolayer to the fraction of NPs that penetrated blank filter inserts. *****P* < 0.0001. (**C**) Penetration of PS 80 (H)-NPs through the cell monolayer in the absence of serum or in the presence of anti-LRP1. **P* < 0.05, ***P* < 0.01. Data in b,c are mean ± SD of technical repeats (*n* = 3, experiment performed at least twice). P values were determined by two-tailed Student’s t-test.

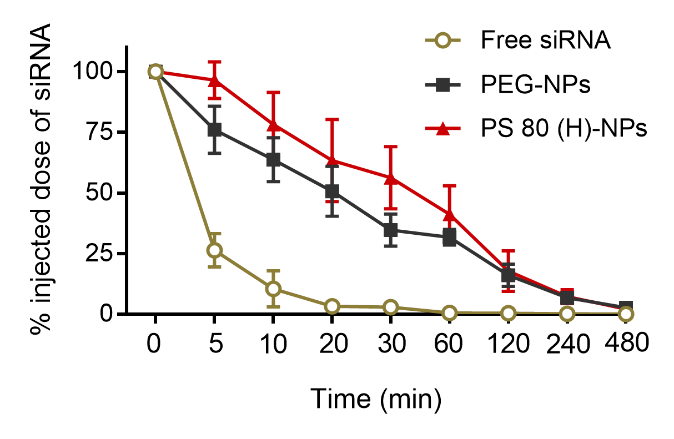

**Fig. S10. Blood circulation profile of siRNA formulations.** Blood circulation profile of Dy677 labeled scrambled siRNA (free), Dy677 siRNA-loaded PEG-NPs, or Dy677 siRNA-loaded PS 80 (H)-NPs after intravenous injection in healthy mice at siRNA dose of 50 nmol/kg. Blood was withdrawn at pre-determined time points and Dy677 fluorescence was quantified. Data in are mean ± SD (*n* = 3 mice/group, experiment performed at least twice).

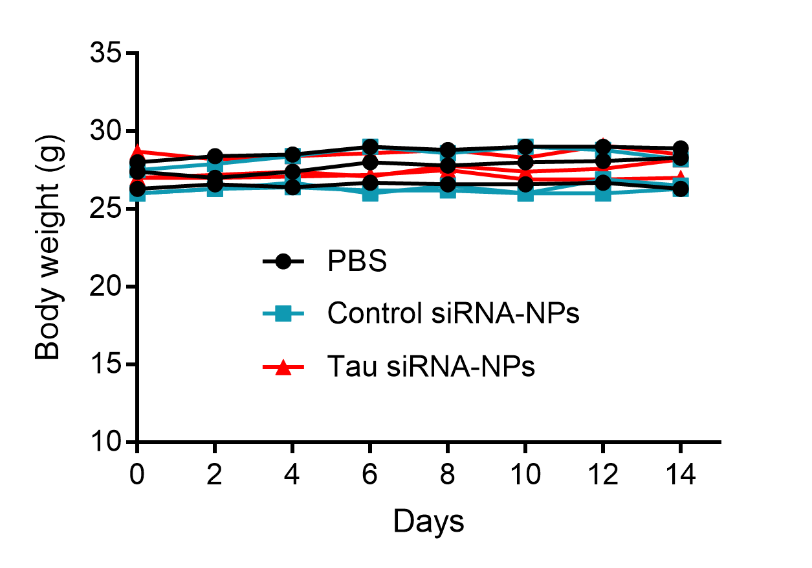

**Fig. S11. Body weight of mice was monitored for assessing the *in vivo* to toxicity of siRNA NPs.** Body weights over time for mice injected with PBS, scrambled (control) siRNA-loaded PS 80 (H)-NPs, or tau siRNA-loaded PS 80 (H)-NPs. Injections were done on day 0 and 1 at siRNA dose of 75 nmol/kg. Data in are mean ± SD (*n* = 3 mice/group, experiment performed at least twice).

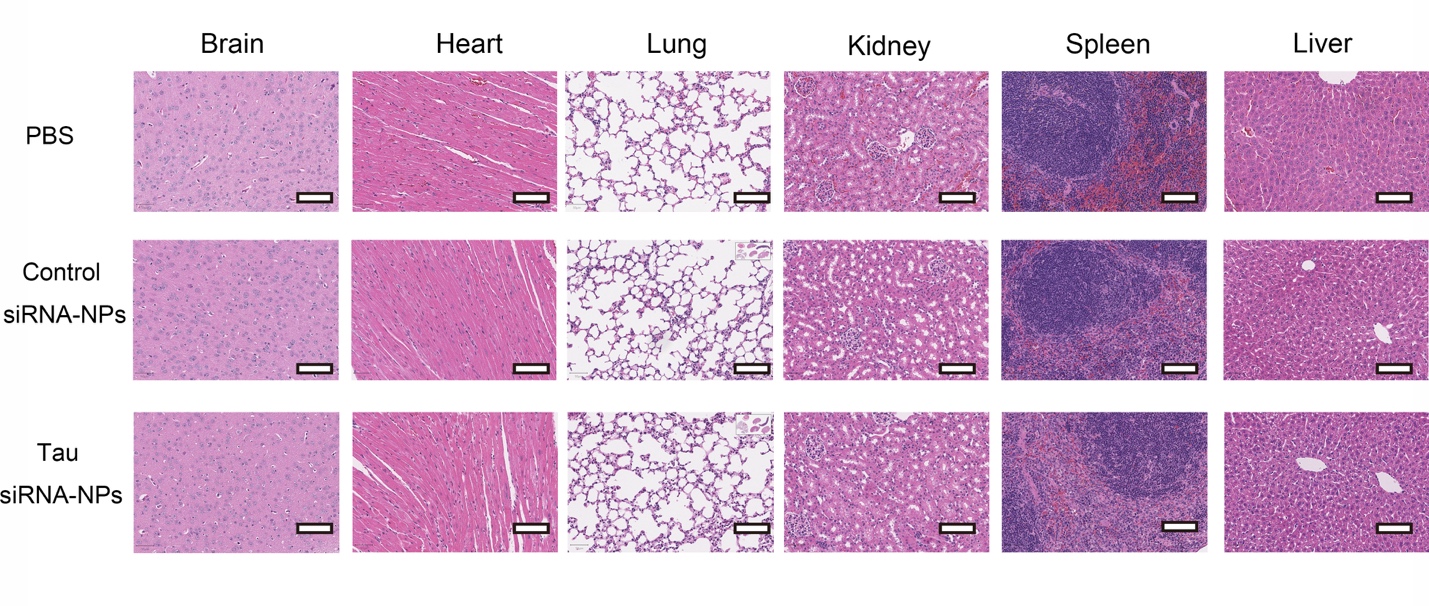

**Fig. S12. *In vivo* safety of siRNA NPs assessed by histopathological analysis.** Representative hematoxylin and eosin (H&E) staining of tissue sections from major organs of mice injected with PBS, scrambled (control) siRNA-loaded PS 80 (H)-NPs or tau siRNA-loaded PS 80 (H)-NPs at siRNA dose of 75 nmol/kg/day. Injections were performed on 2 consecutive days and mice were euthanized 3 days after the last dose. Scale bar = 100 μm.

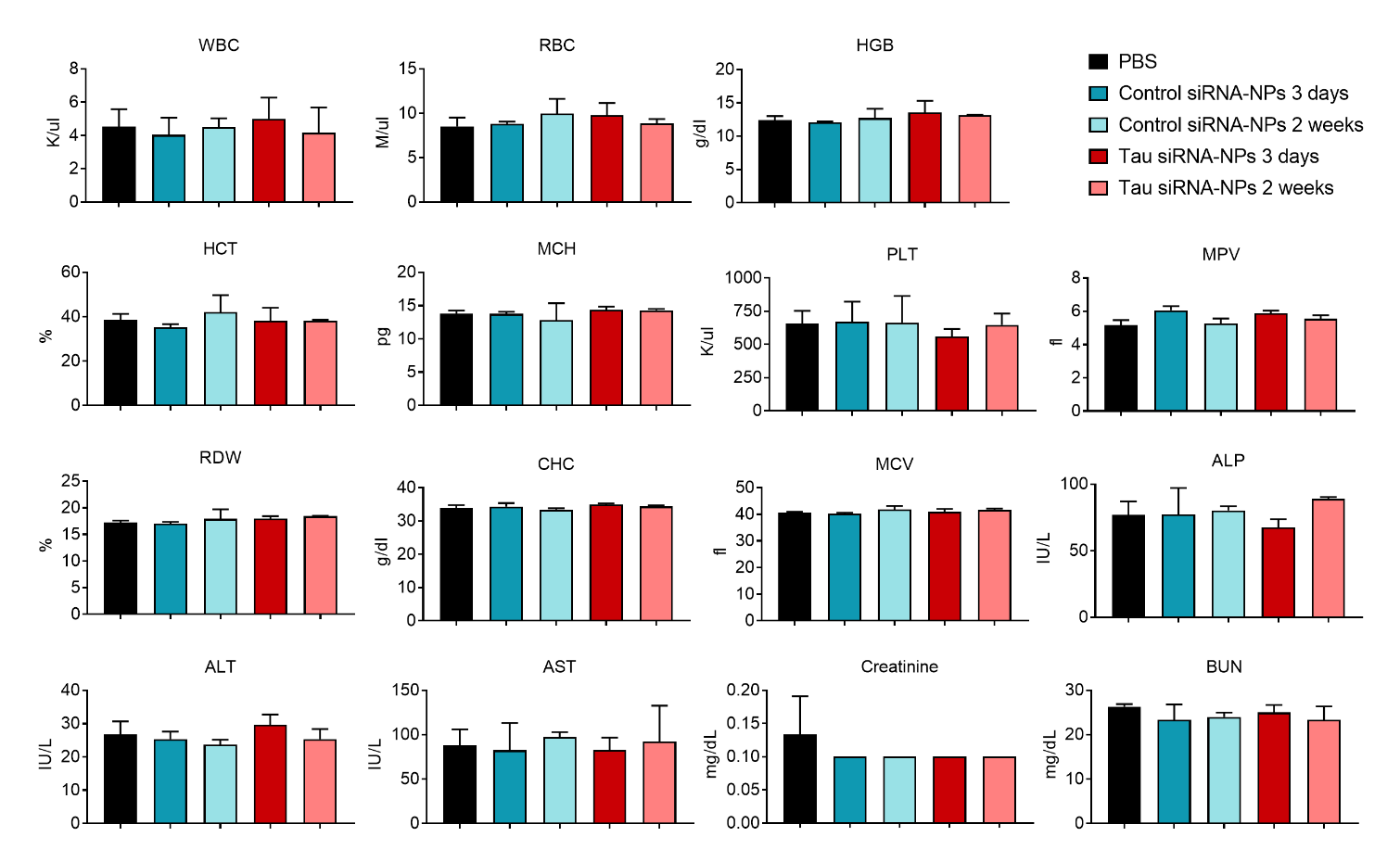

**Fig. S13. *In vivo* safety of siRNA NPs assessed by hematology and blood biochemical parameters.** Hematology and blood biochemical parameters of mice at day 3 or 2 weeks after intravenous administration of PBS, scrambled (control) siRNA-loaded PS 80 (H)-NPs or tau siRNA-loaded PS 80 (H)-NPs. Injections were performed on 2 consecutive days at siRNA dose of 75 nmol/kg/day. Data are mean ± SD (*n* = 3 mice/group, experiment performed at least twice).

**Supplementary Tables**

**Table S1. Coating density of PS 80 and GSH on the surface of NPs.** Total surface area of NPs was determined by assuming spherical shape and uniform size distribution and density for all NPs. Coating density of PS 80 on the surface of NPs was estimated by dissolving NPs in DMSO and subsequently subjecting the solution to HPLC-ELSD analysis. GSH coating density was estimated by assuming 100% incorporation of ligand on the surface of the NPs.

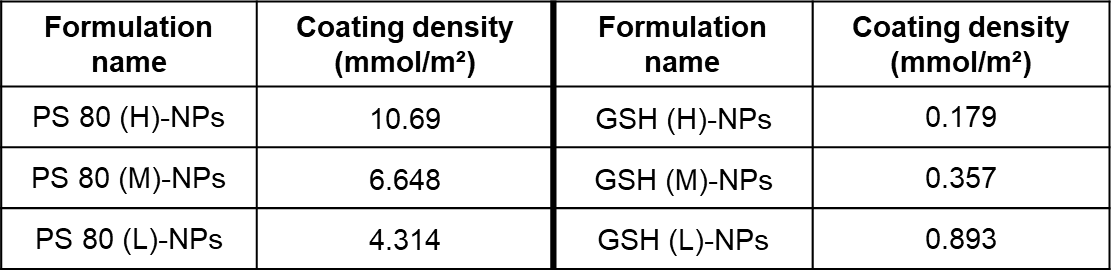
